# Bridging stimulus generalization and representation learning via rational dimensionality reduction

**DOI:** 10.1101/2023.08.09.549352

**Authors:** Lukas Michael Neugebauer, Christian Büchel

**Affiliations:** Department of Systems Neuroscience, University Medical Center Hamburg-Eppendorf, Hamburg, Germany

**Keywords:** generalization, dimensionality reduction, representational learning, Bayesian modeling, fMRI, inductive reasoning, probabilistic inference

## Abstract

Generalization, the transfer of knowledge to novel situations, has been studied in distinct disciplines that focus on different aspects. Here we propose a Bayesian model that assumes an exponential mapping from psychological space to outcome probabilities. This model is applicable to probabilistic reinforcement and integrates representation learning by tracking the relevance of stimulus dimensions. Since the belief state about this mapping is dependent on prior knowledge, we designed three experiments that emphasized this aspect. In all studies, we found behavior to be influenced by prior knowledge in a way that is consistent with the model. In line with the literature on representation learning, we found the representational geometry in the middle frontal gyrus to correspond to the behavioral preference for one over the other stimulus dimension and to be updated as predicted by the model. We interpret these findings as support for a common mechanism of generalization.

---

The real world is very complex and the number of variables that describe it is virtually infinite. Nevertheless, humans are generally very capable of interacting with their environment without having to relearn everything from scratch. This crucial adaptive ability is called generalization and given its paramount importance it is not surprising that extensive research effort has been invested into it [1–6]. The fact that the need to deal with the complex and dynamic nature of the real world is a common underlying problem to any behavior led Shepard [1] to propose a law of generalization as the first law of psychology. In contrast to this common denominator, different subfields have developed largely independently and have treated the generalization of associations, actions and rules separately. These applications map onto research on generalization in associative learning, called stimulus generalization [7], research on the generalization of actions in reinforcement learning (RL), called transfer learning [8] or representation learning [9], and research on reasoning under uncertainty, which we will refer to as inductive reasoning [10]. Beyond those endeavours that are directly concerned with generalization, other research has been concerned with behavioral and neural mechanisms that reduce the complexity of sensory inputs and thereby facilitate generalization. Among those are selective attention [11, 12] and low-dimensional neural representations [13, 14] that can be understood as behavioral and neural dimensionality reduction, respectively.

Stimulus generalization, i.e. generalization of associative learning, describes the transfer of learned stimulus outcome contingencies to novel stimuli [7]. The generalization gradient [15] that results from plotting measured responses against the perceptual continuum that stimuli differ on is often assumed to peak at the learned stimulus and to decrease monotonically with perceptual distance [1, 7, 16]. Due to the intricate link between those gradients and discriminability, the explanatory focus has been on perceptual processes [17–22]. Many studies in stimulus generalization have focussed on fear generalization and its implications for anxiety disorders [2, 3, 23–26]. These and other studies have consistently reported positive neural generalization gradients in the frontoparietal network (FPN) and the salience network (SN) and negative gradients in the default mode network (DMN) [2].

A second line of research that was initiated by Shepard [1] is concerned with inductive reasoning, i.e. the ability to generalize from a set of observations to a general rule. Research on inductive reasoning has produced a series of Bayesian models that are based on a rational analysis of the problem of generalization [27]. These models have emphasized the importance of psychological spaces and conceptualize generalization as the discovery of *natural kinds* which map onto consequential regions in those spaces [1, 10, 28–31].

In contrast to associative learning and inductive reasoning, RL includes actions and a decision making component [32]. An important caveat of the typically used models in the context of RL research in cognitive neuroscience [33, 34] is that without modifications, each state-action pair needs to be encountered multiple times for a reliable estimate of its value and there is no generalization between states. Even in moderately complex tasks, the number of state-action pairs can quickly become intractable [32] and generalization becomes a necessity. A common theme that has emerged in this context is the idea of generalization via abstraction of state spaces. While some studies have suggested the use of correlations in state spaces as mechanism [35, 36], most research focusses on representation learning [9]. This process describes the discovery of an abstracted representation and emphasizes dimensionality reduction [9, 37, 38] or a rescaling of dimensions [39] of the state space. Neural results in this line of research have emphasized the role of the FPN [9, 37, 38, 40], the SN [40–42] and the DMN [9, 43] in the discovery and encoding of abstracted representations.

The idea of generalization via abstraction is related to the dimensionality of neural codes, which refers to the minimum number of spatial dimensions that are needed to account for the variance in neural activity with respect to a certain input space [13, 14, 44–46]. For instance, if stimuli differ on shape and color, only encoding the shape is a low-dimensional encoding and allows for generalization towards new stimuli in tasks where only the shape is relevant. Interestingly, low-dimensional representations have been linked to areas of the DMN [14, 44] and the FPN [45, 46].

Here, we investigated the possibility that these different processes rely at least partly on a common mechanism. For this purpose, we propose a Bayesian model that integrates dimensionality reduction and conducted three studies to test the predictions.

Our model draws on other Bayesian models of generalization [1, 28, 47], but accounts for probabilistic reinforcement and emphasizes the role of the rescaling of dimensions in psychological space according to their perceived relevance for the task. The model assumes *associative maps* [47], which are a mapping from psychological space to the probability of an outcome. Associative maps are characterized by a mean and the strength of exponential decay along each dimension and by the inferred outcome probability at the mean of the map. Since the strength of generalization depends on the weighted distance between novel stimuli and the inferred mean of the map, changing the belief state about the speed of exponential decay along a dimension is exactly equivalent to rescaling the dimension. This approach therefore includes partial dimensionality reduction for small values of the decay parameter and full dimensionality reduction as this parameter approaches 0.

To test how well the model describes human behavior and to explore the neural implementation of dimensionality reduction in the context of stimulus generalization, we designed a series of experiments in which we expected prior knowledge to have a strong influence on behavior. To maximize the role of prior knowledge, we used computer-generated faces that differed on 2 dimensions, identity and emotional expression, as stimuli. We expected emotional expression to carry more information about aversive as well as appetitive outcomes than identity [48]. To distinguish an effect of prior knowledge from an effect of salience only, we manipulated the valence of emotional expression and outcomes. In particular, we used 2 different stimuli sets with either angry of happy faces (fig. 1A-B) and either an aversive (studies 1 and 2) or appetitive (study 3) outcome. Based on the congruency between emotions and outcomes, we expected angry expressions to be associated with aversive outcomes and happy expressions to be associated with appetitive outcomes a priori [49]. Subjects learned to associate one face with an outcome in a conditioning paradigm. The center stimulus (black circles in fig. 1A-B) was reinforced in 50% of trials, while the other faces were never re-inforced. Faces were presented in microblocks in a pseudo-randomized order (fig. 1C) and we collected time-resolved ratings of the outcome expectancy to characterize the dynamics of learning (fig. 1D).

**Figure 1:**
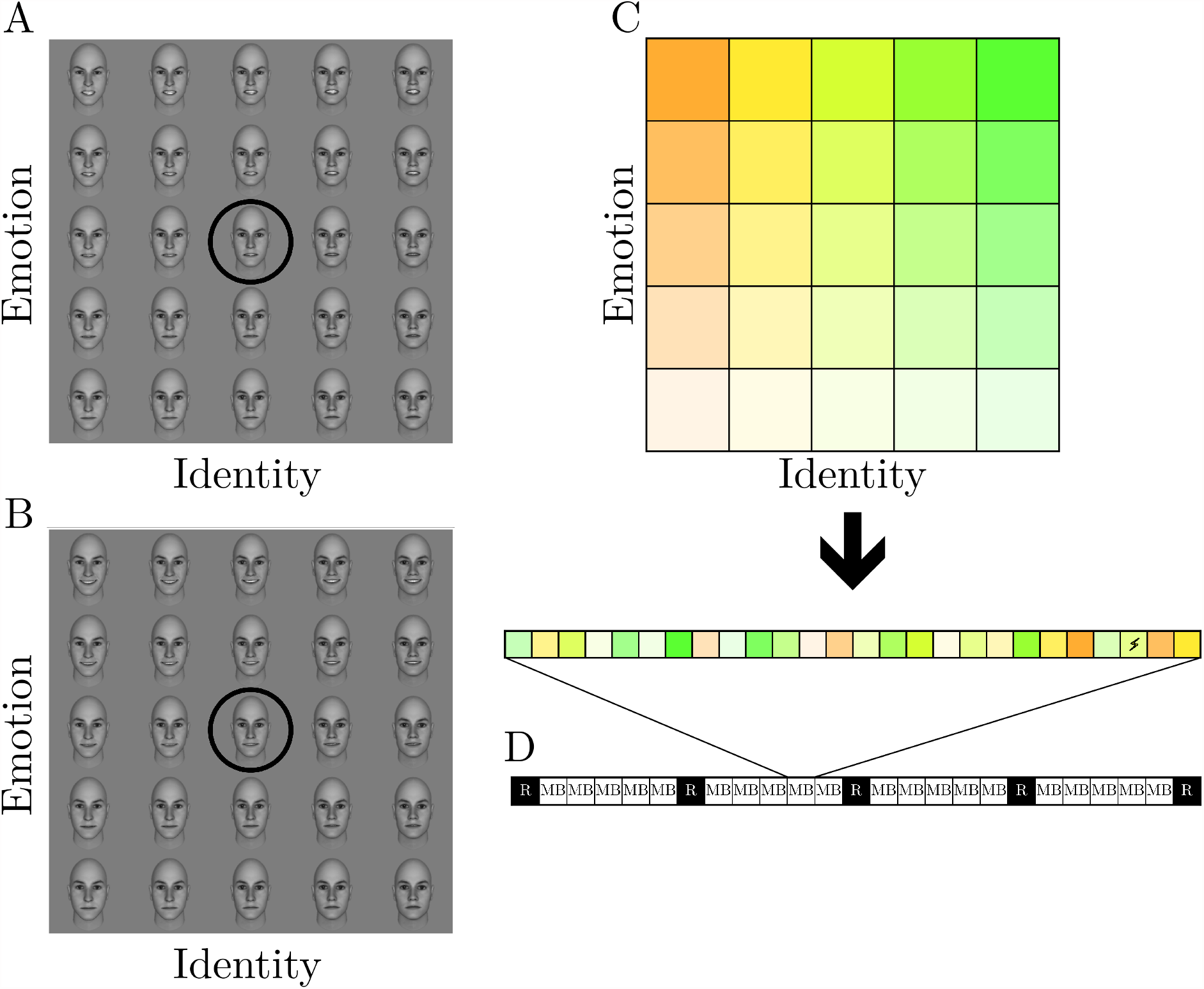
Experimental Design. Stimuli consisted of two 5*x*5 grids of computer generated faces that differed on identity and either (**A**) angry or (**B**) happy expression. The center stimulus served as conditioned stimulus (CS+) (black circle). (**C**) One microblock consists of all faces in pseudorandom order. In addition, either a reinforced trial with the conditioned stimulus (CS+) (75% of microblocks) or an oddball trial (25%) was included. (**D**) The experiment consisted of 20 microblocks (MB), chunked into 4 mesoblocks each. Shock expectation ratings (R) were collected before the first and after each mesoblock for a total of 5 ratings.

In order to make predictions based on the model, we encapsulated the assumed prior knowledge of subjects into appropriate priors for the model parameters and fed the sequence of stimuli and outcomes into the model. Note that these predictions include a prior on the relevance of dimensions and therefore partial dimensionality reduction. From the predictions of our model for this design (depicted in fig. 2A-B), we extracted the following hypotheses: (i) Ratings should initially be primarily dependent on the emotional expression of faces with the direction depending on congruency of the valence of emotion and reward. That is, we expected angry faces to be more strongly associated with an aversive outcome and happy faces to be more strongly associated with an appetitive outcome a priori. (ii) With increasing information (reinforcement), ratings should increasingly depend on the perceptual similarity of stimuli to the reinforced stimulus, (iii) with a differential effect along both dimensions. (iv) The width of generalization and (v) the updating from rating to rating should decrease over time. The extreme case of the differential effect of similarity to the conditioned stimulus (CS+) would be a total neglect of the identity dimension, with learning happening only along the emotion dimension (fig. 2C-D). We include this case because it constitutes a notion of dimensionality reduction that is incompatible with our model. Note that these hypotheses are fairly generic and not exclusive to our model.

**Figure 2:**
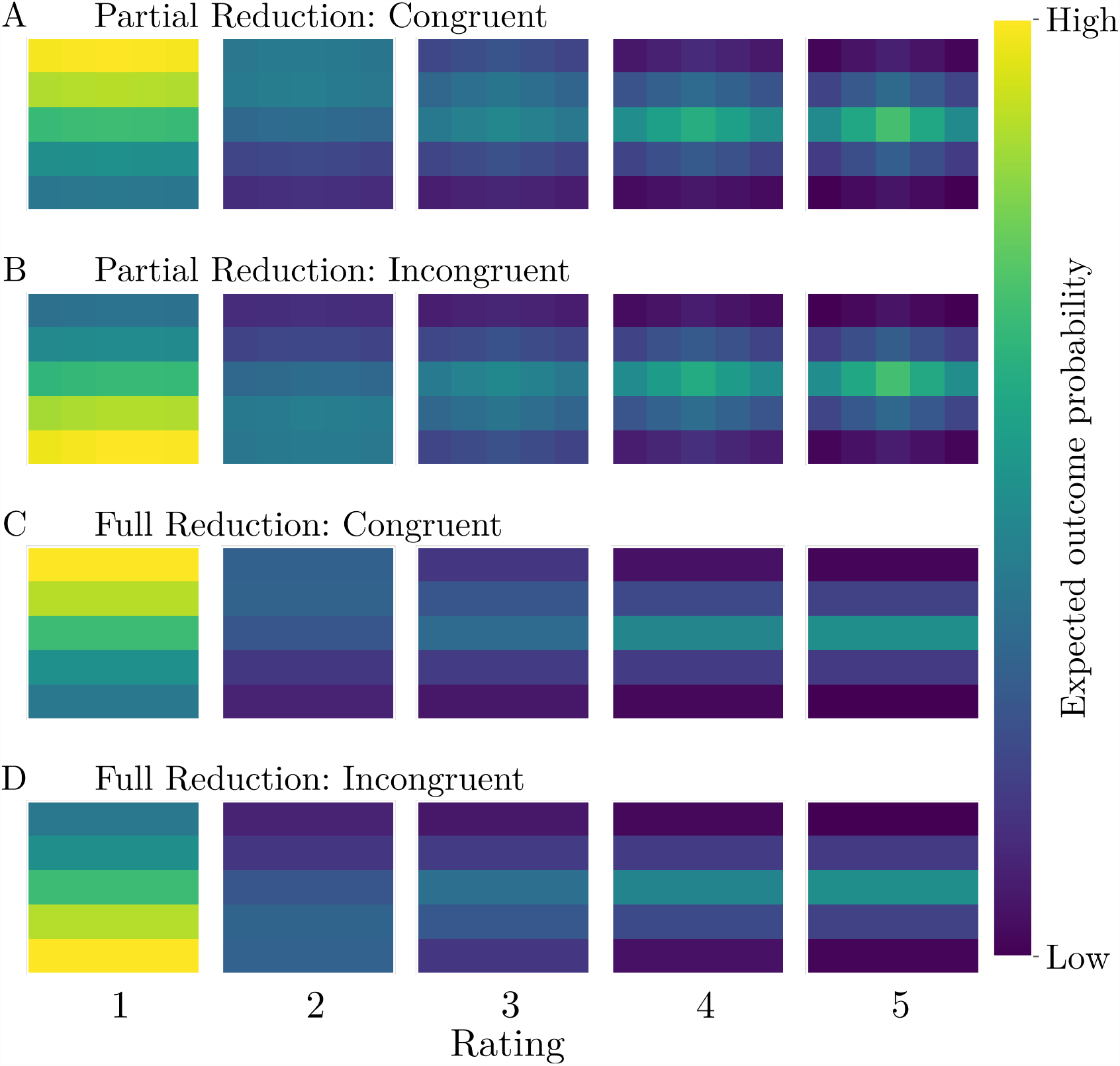
Model predictions. (**A**) Posterior means of model predicted shock expectations for condi tions with congruent emotions and outcomes, i.e. angry faces and an aversive outcome and happy faces and an appetitive outcome. This model assumes a prior about the relevance of dimensions and therefore partial dimensionality reductions. (**B**) The predictions for the same model but incongruent conditions, i.e. happy faces and aversive outcomes and angry faces and appetitive outcomes. (**C-D**) Alternative predictions for congruent and incongruent conditions. These use a model that completely ignores the identity dimensions and therefore encapsulates complete dimensionality reduction.

Behavioral results in all three studies confirmed the predictions of our model, independently of the valence of stimuli and reinforcers. Neurally, we found the middle frontal gyrus (MFG) to encode a scaled representation of the stimulus space using functional magnetic resonance imaging (fMRI). Overall, our results are in line with a common mechanism in generalization.

## Results

### Behavioral results

#### Experiment 1

In the first experiment (N=50, within-subject) we investigated whether shock expectation ratings of subjects followed the predicted pattern in the context of aversive conditioning using a painful electric shock as outcome. To control for differences in perceptual spaces and a potential changes thereof due to conditioning [17–19], subjects completed a perceptual task before and after conditioning. We fit two models to the data from this task that assumed either constant perceptual spaces or spaces that changed due to the learning experience. In both conditions, the model with constant perceptual spaces was favored by the data (tab. S1). Group level solutions of this model suggested that perceptual spaces were well aligned with the intended stimulus space (fig. S1). We included individually fitted perceptual spaces into all subsequent analyses. Because the full Bayesian model is intractable and therefore cannot be fit to data, we used heuristic approximations to the model that included different levels of dimensionality reduction. We fit these approximations in a hierarchical Bayesian manner to the data to further quantify behavior and formally test the model predictions. This is possible because the parameters of these approximated models can directly be interpreted with respect to the hypotheses that we derived from the full model. A visual inspection of averaged ratings for the five shock expectation ratings in the angry condition indicated that subjects’ behavior followed the model predictions quite closely (fig. 3A). We found the model that assumed partial dimensionality reduction to be strongly favored by model comparison (tab. S2). Posterior predictive checks for this model indicated that it captured the observed behavior well (fig. 3A). We further investigated the posterior distributions of this model (fig. 3B). As expected, ratings were initially driven by the amount of emotional expression in faces but increasingly depended on the perceptual distance to the reinforced stimulus. Importantly, this effect was much stronger along the emotion dimension than along the identity dimension. This is also visible in the impact of the different features on the ratings (fig. 3C). The rate of updating decreased and generalization around the center (reinforced) face became narrower over time.

**Figure 3:**
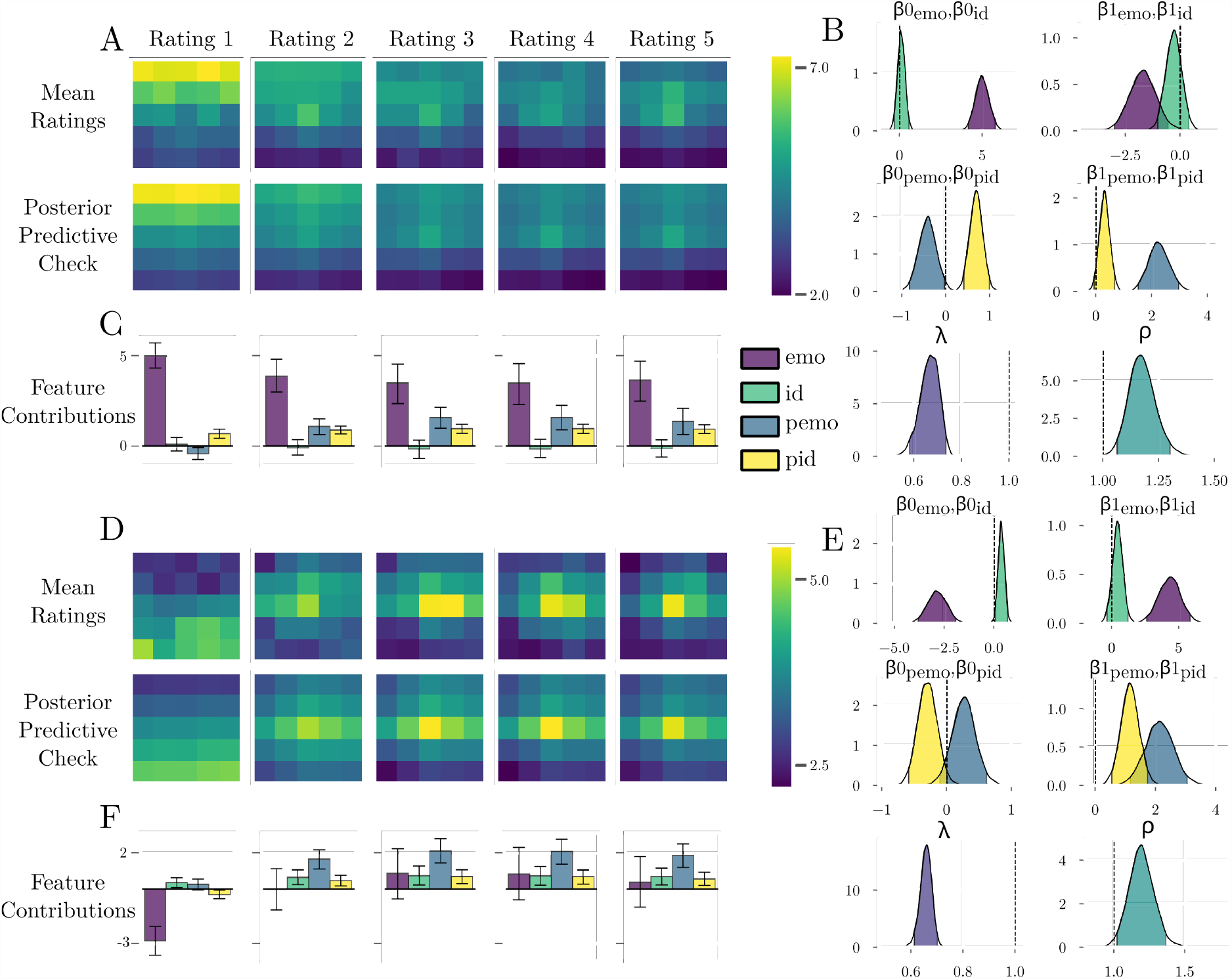
Behavioral results of the first experiment. (**A**) Mean ratings in the angry condition closely followed the predictions from the Bayesian model. A visual inspection of the posterior predictive check indicated that these ratings could be well explained by the fitted model. (**B**) Posteriors of the parameters of the fitted model. Ratings were initially most strongly driven by the amount of emotional expression. We attribute slight deviations from 0 for the initial impact of proximity to the CS+ to a lack of model flexibility. The impact of emotion decreased over time, while the impact of proximity increased, especially along the emotion dimension. The strength of updating and the width of generalization around the CS+ decreased. (**C**) These effects are also visible in the impact of the different features in each rating. (**D**) Mean ratings in the happy condition followed our predictions and could be reproduced with the fitted model. (**E**) The initial impact of emotion showed the opposite sign and was weaker than in the angry condition. In line with the weaker prior, the impact of the proximity along the identity dimension had a steeper slope. Apart from that, the posterior estimates showed the same pattern as in the angry condition. (**F**) The effect of these parameters can be seen in the impact of the different features at each time point.

In the happy condition we found the expected flipped pattern in which subjects rated neutral faces as more dangerous than happy faces in the beginning of the experiment (fig. 3D). For this condition the same model was favored by model comparison (tab. S2) and mean ratings could be closely reproduced by the model as indicated by the posterior predictive check (fig. 3D). Apart from the flipped initial effect of emotionality, the pattern of posterior distributions was very similar to the angry condition (fig. 3E). One interesting aspect was that the initial effect of emotionality was weaker than in the angry condition and that the impact of proximity along the identity dimension had a steeper slope, indicating a weaker prior and an effect of congruency between the valence of the stimuli and the reinforcer (fig. 3F).

#### Experiment 2

In the second experiment we aimed to replicate the findings from the first experiment and to investigate the underlying neural mechanisms. For this reason, we repeated the same design in a new independent sample (N=50, within-subject) while recording fMRI data. To improve the readability of the manuscript, we report fMRI results after the behavioral results of all studies. Using the same model comparison procedure as in the first experiment, we found the model that assumes constant perceptual spaces to outperform the model with dynamic spaces (tab. S2). We again included individually fitted perceptual spaces into all subsequent analyses. With respect to shock expectations, mean ratings showed the same pattern as in the first experiment, with an initial effect of emotionality and a subsequently increasing effect of proximity to the CS+ in both conditions (fig. 4A,D). Using model comparison we confirmed that the model with partial dimensionality reduction was favored by the data (tab. S2) and that the model could reproduce the observed behavior (fig. 4A,D). An inspection of posterior estimates of the winning model and the implied impact of different features at all time points revealed a strong resemblance to the results of the first experiment (fig. 4B,C,E,F) with the only notable difference being faster updating in the scanner. This replication included the stronger impact of emotionality in the angry condition and the flipped initial effect of emotionality in the happy condition.

**Figure 4:**
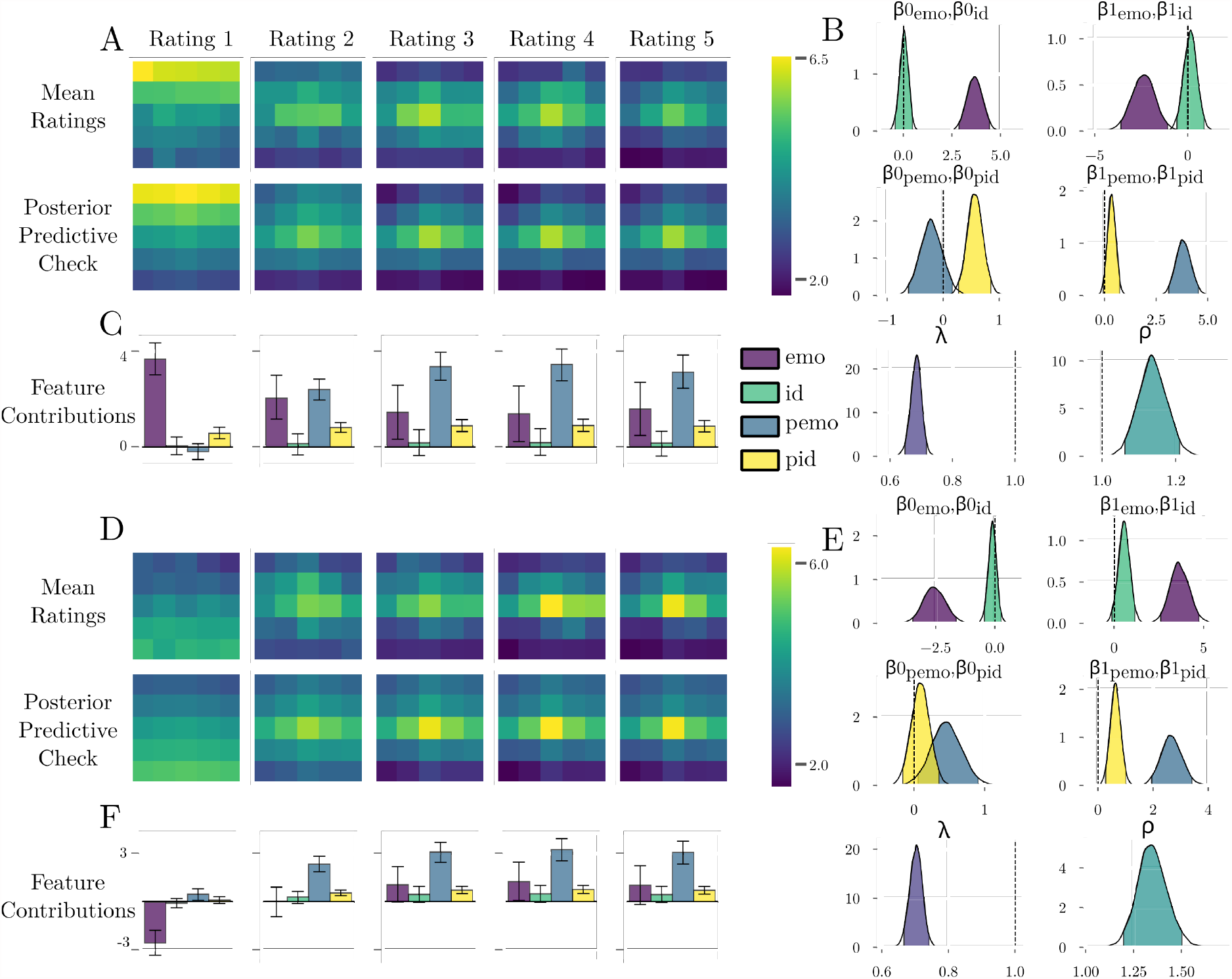
Behavioral results of the second experiment. (**A**) Apart from faster learning, the behavioral results closely mirrored those from the first study and the predictions. (**B**) In line with that, the posterior distributions on model parameters showed the same pattern as in the first study and corroborated our hypotheses. (**C**) The differences of speed of learning can be seen in a quicker dependence of ratings on the proximity along the emotion dimension. (**D**) In the happy condition, we closely replicated the results from the first study. This is also visible in the (**E**) posterior distributions on model parameters and (**F**) the importance of each feature over time.

#### Experiment 3

Since our previous results are based on aversive conditioning, we conducted a third study in an online sample (N=104, between-subjects), where we replaced the painful electric shock with an appetitive monetary reward. This allowed us to broaden the scope of the proposed model and to question the assumption that fear generalization is fundamentally different from other forms of generalization [2, 3]. Due to time constraints, subjects in this experiment completed the perceptual task only once. For this reason, we only fitted a single model to the data. Group level spaces were very similar to the other studies (fig. S1) and we again included individual perceptual spaces in the subsequent analysis. Other than the change in the valence of the outcome and some minor changes that were unavoidable in the context of an online study, the experimental design was otherwise identical to the previous experiments. Note that because we suggest that the initial dependence on prior knowledge depends on the valence of both the stimuli and the outcomes, we expected a flipped pattern of results. That is, we expected happy faces to be considered more and angry faces to be considered less likely to lead to a reward a priori. An initial inspection of mean ratings in the angry (fig. 5A) and the happy condition (fig. 5D) confirmed this expectation. We fit the same models as in the other experiments to the data to investigate this further. Model comparison favored the same model as in the previous experiments in both conditions (tab. S2). The posterior estimates corroborated our initial impression: the results from the first two studies could be closely replicated with the only exception that the initial influence of prior knowledge had the opposite sign: Angry faces were considered to be less likely to be rewarded than neutral faces (fig. 5B-C), while happy faces were rated as more likely linked to an appetitive outcome (fig. 5E-F). Like-wise, while in the aversive generalization studies, the affect of emotionality was stronger in the angry condition, in this appetitive study it was stronger in the happy condition, further corroborating that the congruency of stimuli and outcomes not only affects the direction of the effect but also the magnitude.

**Figure 5:**
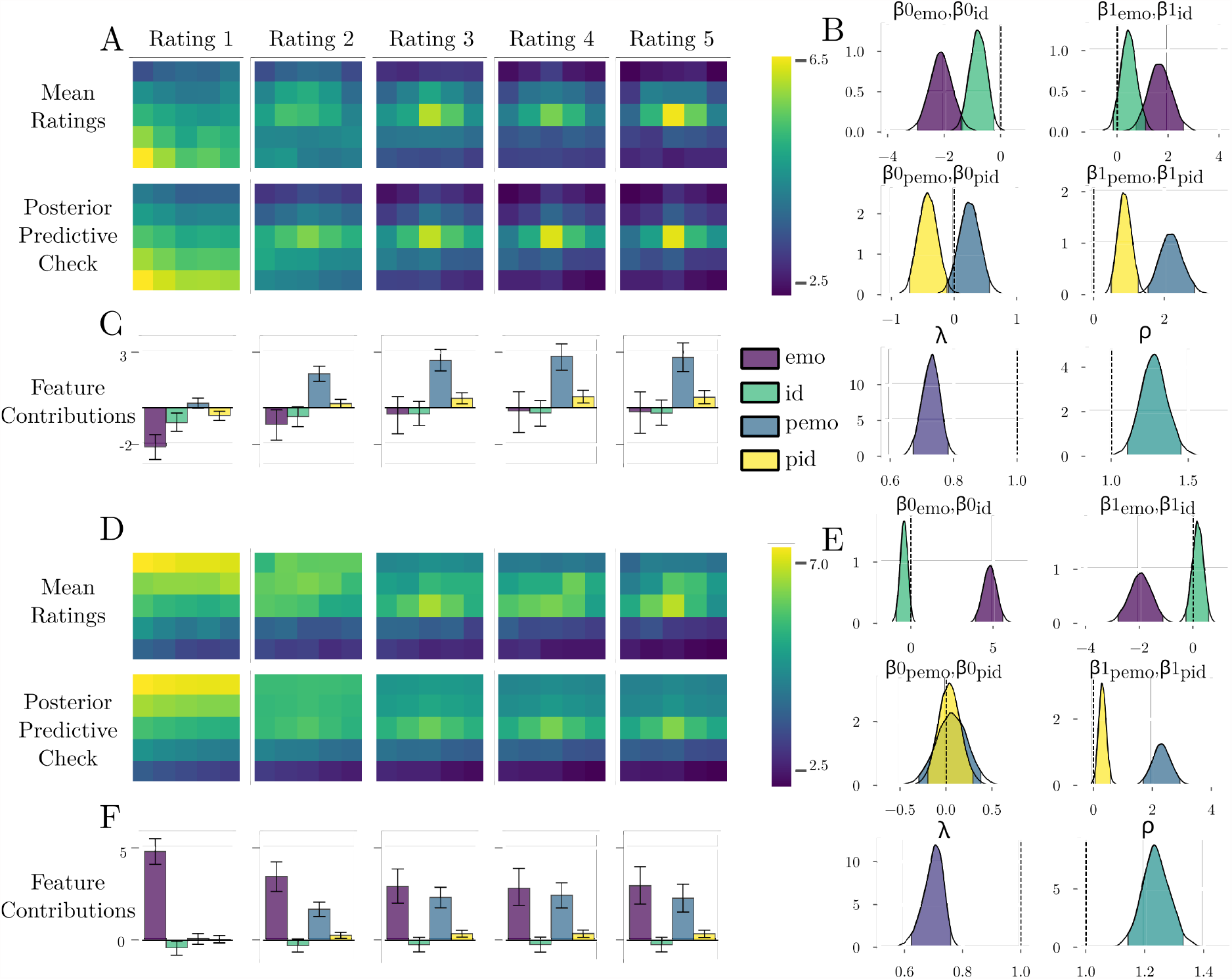
Behavioral results of the third experiment. (**A**) We found the expected flipped pattern of results using appetitive conditioning. Subjects rated angry faces to be *less* likely to lead to a reward than neutral faces. Apart from this, results closely followed the pattern of the previous experiments and our predictions. (**B**) Posterior distributions confirmed this pattern, with the initially strongest effect of emotionality and the increasing effect of proximity to the CS+ that was again stronger along the emotion dimension. (**C**) The impact of features over time is similar to the one in the happy condition of the previous, aversive experiments and therefore matched our predictions. (**D**) Similarly, results from the happy condition are comparable to the angry condition of the previous experiments. This is apparent in the mean ratings, (**E**) the posterior distributions, and (**F**) the impact of features over time. Note that in both conditions we observed an initial unexpected bias towards the left identity. We suspect that this face was perceived as naturally more happy than the right identity.

### Neuroimaging results

We used fMRI data from the second experiment to investigate the neural correlates. In a first step, we interpolated the model-predicted shock expectation to each trial individually and entered those as parametric modulators into a whole-brain general linear model (GLM) (fig. S2). This approach is similar to the typical approach in fear generalization studies [2], but it has the advantage that it tracks the temporal evolution of generalization gradients instead of only the resulting generalization after learning. Nevertheless, directed contrasts revealed the expected pattern of brain activity: we observed positive tunings in the FPN and SN and negative tunings throughout the complete DMN (fig. 6A-C).

**Figure 6:**
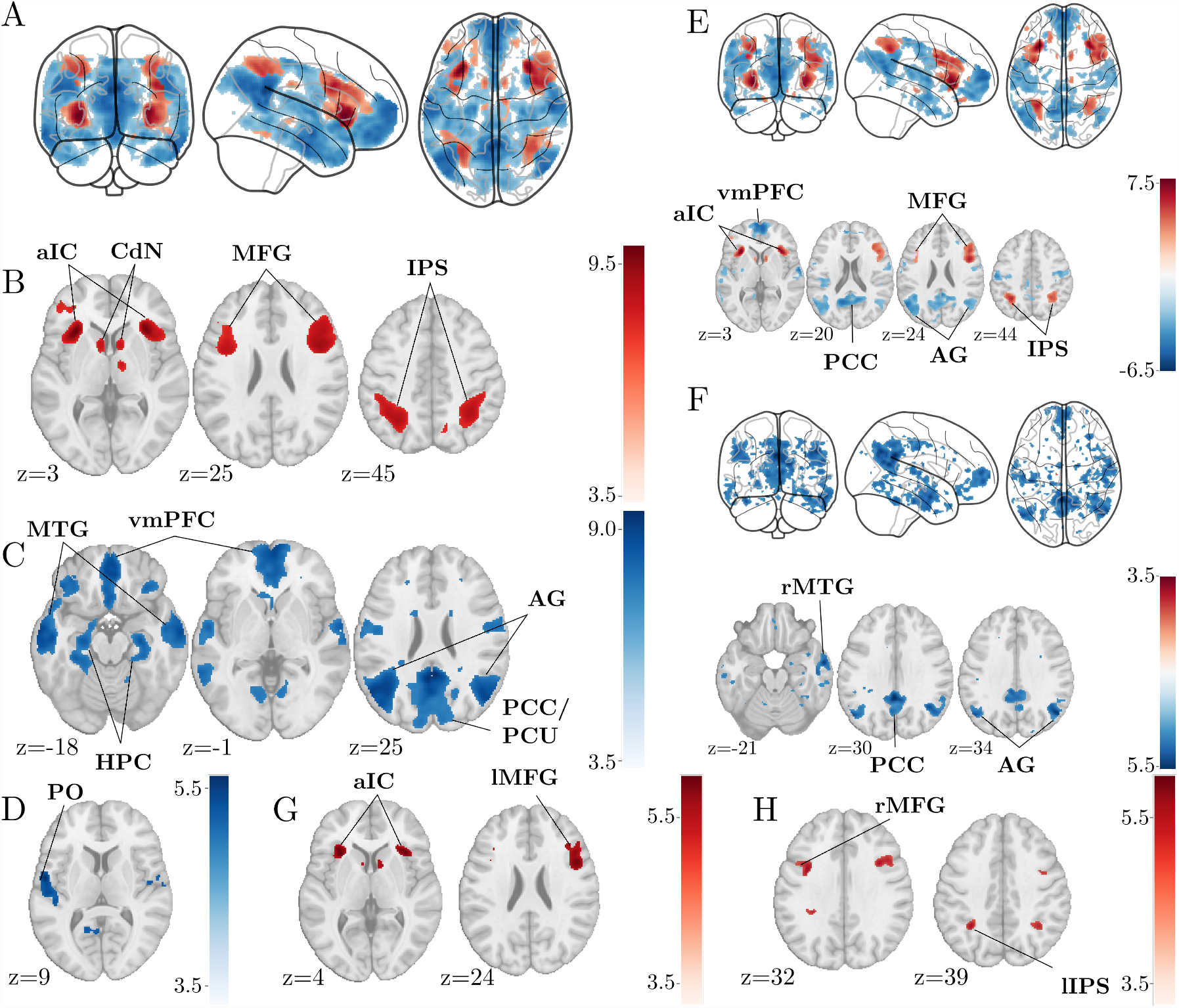
Univariate fMRI results. (**A**) Correlations of interpolated model-fitted shock expectation ratings revealed two distinct networks with different tunings. (**B**) We observed positive tunings in the FPN and SN and (**C**) negative tunings throughout the whole DMN. (**D**) The impact of emotionality on ratings was correlated with activity in the left parietal operculum (PO). (**E**) While correlations of the impact of proximity along the emotion dimension closely matched correlations with the overall shock expectation in all three networks, (**F**) we observed only negative correlations in the DMN for the identity dimensions, but no positive correlations in the FPN and SN. (**G**) We statistically confirmed this in the anterior insular cortex (aIC) and the left MFG with a directed contrast (**H**) and two addition clusters using a small-volume correction (SVC). Note. All results are thresholded at p<.001, uncorrected for visualization. Significant clusters are labeled in the figures.

In a second step, we leveraged the fact that the fitted model conceptualized rating behavior as a linear combination of multiple factors and entered these interpolated features jointly into a second analysis (fig. S2). A single significant negative correlation with the impact of emotion was found in the left parietal operculum (PO), which corresponds to the secondary somatosensory cortex (S2) in humans (fig. 6D). This contrast is especially interesting, because it defines the neural correlates of the prior knowledge about the predictive value of emotional expressions. This deactivation is consistent with studies that report a role of the PO in both the processing of emotional facial expressions [50] and in encoding expectations about somatosensory outcomes [51–53]. This combination makes the PO a likely candidate for the encoding of expectations of painful outcomes given emotion-mediated prior knowledge and suggests that this encoding depends of the modality of both cues and outcomes.

We observed an interesting dichotomy in the neural correlates between the impact of proximity along both stimulus dimensions. While negative correlations in the DMN persisted along both dimensions, positive correlations in the FPN and SN could only be observed with the impact of proximity along the emotion dimension even at a liberal statistical threshold (fig. 6E-F). A differential contrast further corroborated this discrepancy in the anterior insular cortex (aIC), MFG and the left intraparietal sulcus (IPS) (fig. 6G-H). Note that proximity to the reinforced stimulus along the emotion dimension was the strongest driver of ratings over the time course of the experiment and therefore highly correlated with the overall ratings. This can explain the strong overlap between correlations of proximity along this dimension and overall ratings but not the lack of correlations with proximity along the identity dimension in the SN and FPN.

We then went on to investigate the dimensionality of representations more directly using representational similarity analysis (RSA). Because we were interested in the representation of the stimulus space, we constructed representational dissimilarity matrices (RDMs) by computing the absolute distance between stimuli in perceptual space along both dimensions separately (fig. 7A-B). To account for individual differences, we used individually fitted spaces for each participant. Using these model RDMs we performed separate whole-brain search-light RSA analyses for each dimension on beta images that were averaged per stimulus over the complete experiment. Because we reasoned that generalization relies on a representation in which dimensions are rescaled according to their perceived relevance, we computed a differential contrast to identify brain areas where the emotion dimension was more strongly encoded than the identity dimension. In line with our expectations, we found this to be the case in the bilateral MFG (fig. 7C).

**Figure 7:**
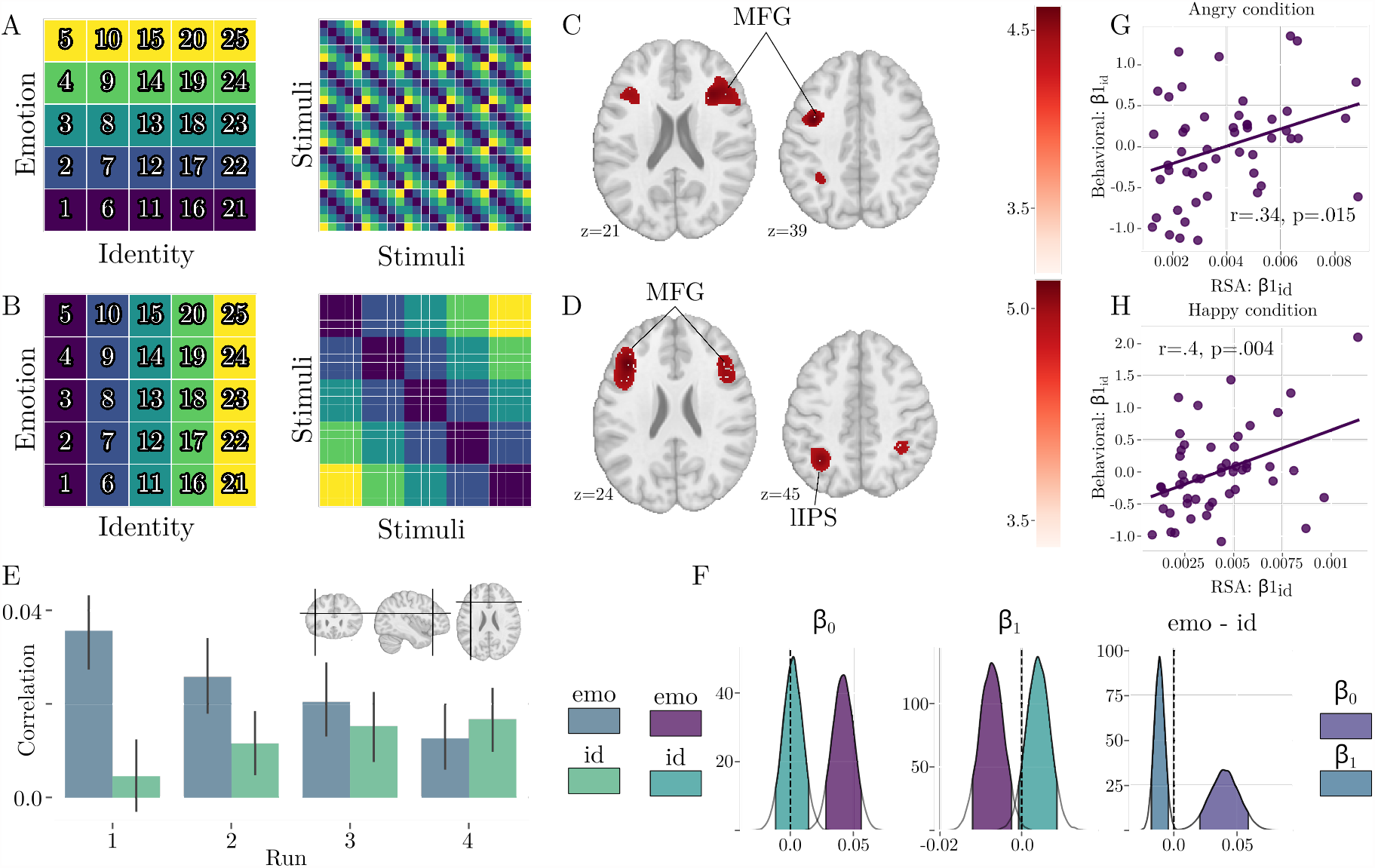
Representational similarity analysis. (**A-B**) We constructed model RDMs by computing the absolute difference of fitted positions along either dimension. (**C**) Neural RDMs in the bilateral MFG showed a stronger correlation with the emotion RDM than the identity RDM when considering the full experiment. (**D**) Using another contrast we identified the left MFG (whole-brain correction) and the right MFG and left IPS as areas in which neural RDMs were correlated with at least one model RDM in at least one mesoblock. (**E**) Representations in the left MFG were initially dependent on only the emotion dimension, but increasingly dependent on both dimensions. Inlay depicts the location of the peak voxel. (**F**) Posterior estimates of a hierarchical linear regression confirmed this pattern. Neural RDMs in the left MFG were initially correlated with the emotion RDM (*p*(*β*0_*emo*_ *>* 0) ≈ 1), but not the identity RDM (*p*(*β*0_*id*_ *>* 0) ≈ .594). The correlation increased for the identity RDM (*p*(*β*1_*id*_ *>* 0) ≈ .919) and decreased for the emotion RDM (*p*(*β*1_*emo*_ *>* 0) ≈ .005). Posteriors on the differences between the dimensions confirmed this for the intercept (*p*(*β*0_*emo*_ − *β*0_*id*_ *>* 0) ≈ .999) and the slope (*p*(*β*1_*id*_ − *β*1_*emo*_ *>* 0) ≈ .997). (**G-H)** The slope of correlations with the identity RDM in the left MFG was correlated with the increased importance of the identity dimension for ratings in both conditions.

In the context of our model, this discrepancy maps onto the differences in the perceived relevance of dimensions. If the scaling of dimensions in neural representation indeed reflected this belief state, we expected it to be updated analogously. To investigate this, we repeated the RSA searchlight, but used beta images that were averaged per fMRI run instead of the whole experiment, resulting in four searchlights per subject and dimension. Before characterizing the evolution of representations, we used a generic F-contrast to identify areas in which the representation depended on at least one dimension in at least one run while averaging over conditions. This contrast revealed a cluster in the left MFG and two more clusters in the right MFG and the left IPS when using small-volume correction (SVC) in a mask of the FPN [54] (fig. 7D). We then extracted the correlations between the model RDMs and the neural RDMs per run from the peak voxels of these clusters. In line with their expected correspondence to the Bayesian model, correlations in the left MFG showed a pattern in which representations initially depended exclusively on the emotion dimension and then gradually evolved to depend on both dimensions (fig. 7E). While a similar pattern could be observed in the right MFG, representations in the left IPS were more static and depended on both dimensions throughout the experiment (fig. S3). To investigate these patterns more formally, we modeled the time course of correlations using a Bayesian hierarchical linear regression. While we refrain from interpreting posterior estimates in a frequentist manner, the bulk of probability mass in these estimates corroborates the pattern of results described above (fig. 7F). Since we stipulated a correspondence between those representations and the observed behavior, we expected a statistical relationship between the neural and the behavioral data. In particular, we reasoned that the increase in the relevance of the identity dimension in behavior should be reflected in an increased dependence of neural representations on this dimension. We therefore correlated the posterior estimates of the change in relevance of proximity along the identity dimension in behavior with the slope of the change in the relevance of the identity dimension in the left MFG and found significant positive correlations in both the angry and the happy condition (fig. 7G-H).

## Discussion

Based on a consideration of how general the problem of changing contexts affects any real world agent, we aimed to explore the possibility of a mechanism of generalization that is common to the many applications that have been proposed in the literature. In an attempt to propose an integrative framework, we have therefore formulated a Bayesian model that comprises aspects from the literature on stimulus generalization, representation learning and inductive reasoning. Since the predictions of this model strongly depend on the assumed prior beliefs, we conducted three studies that emphasize this aspect.

All three studies strongly indicate that subjects’ behavior closely followed the predictions of our model. In particular, expectation ratings initially depended on the prior knowledge of subjects as they were almost exclusively driven by the amount of emotional expression. Importantly, this was the case for appetitive and aversive conditioning in a congruent manner: Angry faces were rated as more likely to lead to an aversive outcome while happy faces were taken to be more predictive of an appetitive outcome. This preference for the emotion dimension extended to faster learning about this dimension despite the same amount of exposure to both dimensions. Neurally, we observed an interesting dichotomy in the usually implied networks involved in fear generalization with a strong preference for the relevant dimension in the aIC and the MFG. Finally, we report a correspondence of the evolution of neural representations in the left MFG with the dynamic relevance of the dimensions in behavior and a correlation between the two.

Our work makes a number of important empirical and conceptual contributions to the literature on generalization. With respect to fear generalization, our results strongly challenge a number of widely held assumptions. First, we conducted studies using either appetitive or aversive conditioning and found strong parallels between them. More specifically, the mean ratings and parameter estimates indicate that both appetitive and aversive generalization rely on the same mechanism when accounting for the congruency between the valence of the outcome and the valence of the emotional expression. While some literature suggests specific mechanisms in fear generalization with respect to perceptual tunings [17–19], the strong overlap in our results agrees with studies that have investigated appetitive generalization and reported generalization gradients that closely resembled those in fear generalization [55–57]. Beyond merely challenging the idea of fear generalization as a distinct process, our model provides an explanation for the increased fear generalization in the context of anxiety disorders [23, 24], which is likely the reason for this assumption. In particular, we propose that the increased fear generalization in anxiety disorders is a consequence of maladaptive prior assumptions about the shape of associative maps or – to put it in the context of Shepard [1] – the size of consequential regions.

Second, we challenge assumptions about the role of perception in stimulus generalization. The most extreme case of this is the idea that generalization is fundamentally a failure to discriminate between stimuli [5]. Although this mechanism lacks face validity and parsimony, as it would require different mechanisms for observations like peak shifts [7, 58] and monotonic generalization gradients in intensity generalization [59], it is still being discussed in the literature [20–22, 60]. Multiple aspects of our data are incompatible with a purely perceptual mechanism. First, we observed a remaining bias towards the emotional expression even after extensive training. Second, we report stronger generalization along the identity dimension than the emotional dimension in all of our studies and conditions, although we carefully controlled for perceptual spaces. However, it is noteworthy that this does not rule out low-level mechanisms that are specific to aversive conditioning like a perceptual retuning around negatively reinforced stimuli [17– 19].

More generally, our results are in line with a mechanism of generalization that comes *before* associative learning or RL. This assumption agrees with Shepard’s universal law, with previous attempts in the literature to integrate subfields and with parallels between them that have largely gone unnoticed so far. A few studies have combined associative learning with instrumental generalization phases in which subjects used the learned associations to guide their behavior [61–63]. While this approach differs from RL because the learning phase is passive, it provides a bridge to work on generalization in RL. These studies reported instrumental generalization gradients that were similar to those found in stimulus generalization [61] and strongly correlated with associative generalization gradients within subjects [62, 63]. In addition, while the vast majority of studies on stimulus generalization have used one-dimensional stimulus spaces and therefore have not explored dimensionality reduction as a mechanism, results from the few studies that have used multidimensional stimuli are well in line with this idea [64–66]. Lee *et al*. [47] explicitly linked inductive reasoning to stimulus generalization by adapting a Bayesian model of inductive reasoning to account for probabilistic reinforcement. They concluded that both phenomena rely on similar processes.

Conceptually, both inductive reasoning and generalization in RL have a spatial component, psychological spaces and state spaces respectively. Consequential regions in psychological space and abstractions of state spaces seem closely related. More explicitly, this parallel can be seen in newer developments. In a Bayesian model by Soto *et al*. [31], consequential regions are parameterized by a mean and the width along each dimension. This shows a close resemblance to very recent developments in RL, where generalization is assumed to rely on the mean and covariance of observed examples in some cognitive space with a rescaling of dimensions according to their perceived relevance for the task [39]. Likewise, exploiting correlations in the reward probability of tasks [35, 36] is conceptually similar to the model of Lee *et al*. [47], where generalization is based on the assumption that similar stimuli lead to similar outcomes. In general, a psychological space is an adequate abstraction to begin with and while Shepard [1] suggested that psychological spaces are shaped by evolution, they are likely to be shaped by learning as well [67].

Our work explicitly links representation learning to stimulus generalization in the context of a model that is based on those proposed in inductive reasoning. This is important since it provides researchers in all three fields with a common language. In addition, it omits the distinction between the learning of an abstract representation and the learning of associations between this representation and outcomes and provides a rational explanation for the observed partial dimensionality reduction.

On a neural level, we provide a mechanistic interpretation of parts of the typically observed neural correlates in fear generalization and link them to the literature on representation learning. Despite some attempts to characterize the role of different brain areas [e.g. 16, 62, 68], most of the neuroimaging literature in fear generalization has been concerned with identifying brain areas that show a generalization tuning [2]. Because of their correlational nature, these analyses can not distinguish between a correlate of learning itself and a correlate of the result of learning. More specifically, the pattern of observed tuned areas is compatible with an interpretation that is purely based on the salience of stimuli after conditioning and a SN mediated switch from a default to a task state [69]. Somewhat agreeing with this, the strong overlap of neural generalization tunings with the three major brain networks has been noted in the literature [70, 71] and included in a new iteration of a neural model of fear generalization [2]. Here, the SN is assumed to be involved in threat processing, the DMN in the reinstatement of a default mode and the FPN in the deployment of attentional resources. Deviating from this, our data shows a clear role of the FPN, especially the MFG, in the encoding of an adequately abstracted representation of the stimulus space. This finding is in agreement with results from representation learning [9, 37, 38] and task representations more generally [12, 40–42, 72–74]. In addition, the continuous updating of this representation fits very well with a recently proposed minimal representation strategy in the lateral prefrontal cortex (PFC) [75] and with research that suggests an ability of the MFG to flexibly switch between low- and high-dimensional representations according to task demands [45, 46, 76]. A noteworthy caveat with the interpretation and generalization of the results stems from disagreements in the literature about representation learning and task representations. While some authors have argued that the FPN is involved in the discovery *and* encoding of representations [40, 73], others agree about the discovery, but locate the actual encoding to the aIC [42] or the orbitofrontal cortex (OFC) [9], which agrees with its role in state space representations and cognitive maps [43, 77, 78]. While our results agree with an actual encoding in the MFG, this remains an open question for future research.

In summary, we have shown that rating behavior in aversive and appetitive generalization closely followed predictions from a model that combines stimulus generalization, representation learning and inductive reasoning. While we are aware of the relatively general nature of our hypotheses, we believe that our model provides a useful framework for future research because there are some natural extensions that can be explored. For example, future iterations could explore other gradient shapes or a distribution over different possible shapes, the inclusion of action spaces and different types and magnitudes of outcomes. In addition, our neural results provide a bridge between the literature on fear generalization and representation learning, task representations and the dimensionality of neural codes. We hope that this work motivates future research in this direction and that it will help to bridge the gap between the different fields.

## Methods

### Ethics approval

All experimental procedures were approved by the Ethics committee of the General Medical Council Hamburg (PV 4164).

### Participants

#### Studies 1 and 2

All participants had normal or corrected-to-normal vision and reported no history of neurological or psychiatric disorders. Written informed consent was obtained before the start of the experiments. Subjects were paid 12 €/h for the approximately 2.5 hours of participation and stated to not have participated in a study including face stimuli and aversive learning before.

For the behavioral sample (study 1) we collected data from 53 subjects. Three subjects dropped out after the first day of the experiment. The remaining 50 subjects were included in the analysis (35 female, mean age=24.96, range 18-37). Due to scheduling conflicts, we could not collect data for the quadruplet tasks (described below) for 1 subject each in the happy and angry condition. For these subjects we exploited the hierarchical structure of our perceptual model and entered the group level perceptual space in the following analysis while we used the subject level perceptual space for all other subjects. We discarded 3 more subjects from the behavioral models in the happy condition, 1 because they were reporting a constant shock expectation of 1 during all ratings and 2 more because their parameters would not converge between MCMC chains 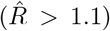. Accordingly, we fit all behavioral models in the happy condition with data from 47 subjects and models in the angry condition with data from 50 subjects.

For the fMRI study (study 2) we collected data from 62 subjects. We discarded a total of 12 subjects. The reason for the relatively high dropout rate is that we only included subjects that had usable fMRI data in both sessions. 2 subjects showed excessive movement, 2 subjects fell asleep (verified through a face camera), 1 subject showed structural abnormalities in the anatomical image, for 4 subjects the electrode for the electrical shock dispatched during data acquisition and 3 subjects dropped out after the first session, leaving us with 50 data sets (27 female, mean age = 26.42, age range = 18-40). Due to scheduling conflicts we omitted the quadruplet tasks for 3 subjects in the angry condition and 2 subjects in the happy condition. We treated these cases as outlined above.

#### Study 3

Participants for the online study (study 3) were acquired using Prolific (https://www.prolific.co). The study itself was hosted on Pavlovia (https://pavlovia.org/) and we obtained written consent using the online interface. We constrained eligible participants to those who reported German as their first language, were between 18 and 60 years of age, had at least completed 50 previous submissions and an approval rate of at least 95%. We collected data from a total of 104 participants (44 female, mean age = 30.92, age rage = 18-59). Subjects were randomly assigned to either the angry or happy condition, which resulted in 48 subjects in the angry and 56 subjects in the happy condition.

#### Stimuli

Face stimuli were created using FaceGen (FaceGen Modeller 2.0, Singular Inversion). We created 2 initial facial identities and subsequently generated 25 faces in a grid-wise fashion by morphing the two identities in 5 steps (100% first identity to 100% second identity) and adding 5 morphing steps of emotional expression (angry and happy each) from no emotional expression to strong expression to each of the neutral faces that resulted from morphing the identities. The individual perceptual spaces were fitted as outlined below.

### Experimental design overview

#### Studies 1 and 2

We employed a two day within-subject design in both experiments. Before starting the experimental procedures on the first day, we acquired informed consent. On both days, subjects initially went through a perceptual quadruplet task (described below) in a behavioral lab which allowed us to estimate their perceptual spaces. The generalization experiment was conducted in a behavioral lab for the first study and in the MRI scanner for the second study. Before starting the actual experiment, we calibrated the electric shock to a painful, but bearable level (see below). After the calibration, stimuli and shocks were presented in a sequence of 20 microblocks chunked into 4 mesoblocks of 5 microblocks each. This approach is based on Onat [79]. We collected shock expectation ratings before, after and in between the mesoblocks for a total of 5 ratings. After the generalization experiment, subjects repeated the perceptual quadruplet task in a behavioral lab to control for changes in perceptual spaces.

#### Study 3

In contrast to the previous studies, we used a between-subjects design due to the added difficulty of running a within-subjects study in the context of online data collection. Eligible subjects chose to participate in our study on Prolific and were redirected to Pavlovia, where they completed their participation. Subjects received 12 £/h for the approximately 105 minutes duration of the experiment plus a bonus of up to 4 £(see below). Instead of an electric shock we used monetary reinforcement and collected reward expectation ratings as the behavioral readout. We also omitted the repetition of the perceptual task after the generalization task to keep data quality high and because we established a constant perceptual space in the two previous studies. Apart from that, the design was equivalent to the previous studies.

#### Quadruplet task

Subjects from all experiments completed a two alternative forced choice task (2-AFC) to generate data that we could use to estimate their perceptual spaces using a hierarchical Bayesian approach that we developed based on maximum likelihood difference scaling (MLDS) [80] (see below). For each trial, subjects were shown a quadruplet of faces, in which the upper two and lower two faces each formed a pair. Subjects were asked to indicate which of the pairs was more similar within-pair and pressed the ‘y’ or ‘m’ key (QWERTZ keyboard layout) to indicate their decision for the upper or lower pair respectively. We used the up and down arrow keys instead in the online study because we could not control the keyboard layout that subjects were using. The response was self-paced. Because there is an unreasonably large number of potential quadruplets for 25 faces and we could not expect to accurately fit the perceptual space with an insufficient number of sampled pairings, we only used the 9 faces that form the outer 3 × 3 grid (including the center stimulus) and interpolated the perceptual space based on the fitted positions of these 9 stimuli (see below). The task was split into 4 blocks of 97 trials with self-paced breaks between them for a total of 388 trials. The number of trials was based on a previously reported study from our lab. In this study, 216 trials were used to estimate the positions of 8 stimuli. To account for the added complexity of estimating 9 positions, we opted to use the same percentage of possible quadruplets (12.8%). We generated an informative sequence of quadruplets by running simulations using a hypothetical observer and 2000 random sequences of quadruplets and took the sequence for which the distances between the assumed and the fitted stimulus positions were minimal. This sequence was used for all subjects in all experiments. This task was repeated before and after the main experiment in studies 1 and 2 in order to control for changes in perceptual spaces. After having established a constant perceptual space, we omitted the second iteration of the task in study 3 to shorten the duration of a study and prevent a decline in data quality.

#### Pain calibration

We set the amplitude of the electric shock to 1.5 times the subjective pain threshold, i.e. the amplitude that is rated as painful on half the trials. To calibrate this pain threshold, we used a QUEST procedure [81] with two alternatives „Painful” and „Not painful”. The calibration consisted of 12 trials with amplitudes that were suggested by the QUEST procedure. The electric shock was delivered by a direct current stimulator (Digitimer Constant Current Stimulator, Digitimer) via an electrode that was connected to the back of the left hand. Shocks consisted of trains of 5 ms pulses interrupted by 10 ms breaks for a total of 100 ms. For the fMRI study we ran a dummy EPI sequence during calibration to control for putative effects of the noise on pain perception. Prior to starting the experiment, the shock intensity was validated to be painful but bearable. No calibration was used in the online study because we did not use electric shocks.

### Main Experiment

#### Studies 1 and 2

For the main experiment, subjects were informed about the procedure via written instructions on the monitor. After that they were presented with 9 random faces and the oddball face to familiarize themselves with the experiment. The oddball consisted of a face that was smoothed with a Gaussian kernel to not be recognizable anymore. Subjects were instructed to press a key as soon as the oddball face appeared to ensure ongoing attention. Familiarization trials were identical to the trials of the microblocks, but subjects were informed that they would not receive a painful shock in this phase. After that we acquired the first shock expectation rating. For this purpose, we presented each face in random order for 1.5 s. After the stimulus offset, subjects should indicate their response to the question „How likely is this face going to be followed by an electric shock?”. They were presented with a 10-point Likert scale with the anchors „not at all likely” (1) and „very likely” (10). Subjects had to select their answer with the left and right key and confirm with the up key. In study 1 those were the arrow keys on the keyboard and in study 2 the left, right and up button on a response button box with a diamond button layout in the scanner. Note that the first rating is before any reinforced trials have been presented in order to probe the prior expectations that subjects brought into the experiment. After that we proceeded with the presentation of stimuli and shocks in a microblock-wise manner. Each microblock consisted of 27 trials. One trial per all 25 stimuli, including a not reinforced trial for the probabilistically reinforced stimulus, one null trial without any stimulus and one reinforced or oddball trial. Per mesoblock of 5 microblocks there were 4 reinforced trials and 1 oddball trial. Stimuli in the main experiment were presented in a truncated type 1 index 1 (t1i1) sequence (37). The reason was to prevent putative higher-order learning as these sequences make it impossible to predict which stimulus is about to follow. For this purpose we ran the designseqran C++ program (retrieved from https://www.bioss.ac.uk/people/cmt/designseq.html) for one hour to generate random t1i1 sequences with N=27 (25 stimuli + 1 reinforced or oddball trial + one null trial). One reinforced trial per mesoblock was replaced with an oddball trial to enforce attention. Because of time constraints that were imposed to prevent declining data quality and the expected speed of learning, we then generated all possible subsequences consisting of 20 microblocks instead of the full 27 iterations of the complete t1i1 sequence for all full sequences. We determined the best sub-sequence with the respect to criterion 4 (37) and generated sequences for individual subjects by keeping reinforced, oddball and null trials and non-reinforced trials of the reinforced stimulus constant and replacing the rest with random permutations of the stimulus indicators. Each trial started with a fixation cross on the monitor that was followed by the presentation of the stimulus after 850 ms. The stimulus was presented for 1.5 s and the fixation cross jumped from the forehead to the mouth region after 750 ms. Subjects were required to follow the fixation cross to ensure that they gathered information about the full face and to prevent an effect of idiosyncrasies in gaze patterns. In reinforced trials, the electric shock occurred 100 ms before stimulus offset. The ITIs were 3, 4, 5 or 6 seconds, each with a probability of 25 %. We made sure that ITIs were equally distributed over the trial types.

#### Study 3

Subjects were informed about the experiment with written text on the monitor. Apart from the fact that this was an online study, the only difference to the previous studies was the use of a different reinforcer. We informed subjects that sometimes a face could hand out a reward. To implement this, we showed a one dollar bill in conjunction with the rewarding face for 2 seconds after the stimulus offset in rewarded trials. Subjects were told that each reward would potentially be worth 25 pence and that they would receive this payment on top of the baseline compensation if they reacted quickly to all oddball trials by pressing the space bar on their keyboard. They were informed that the bonus that they actually received was proportional to the amount of oddball trials that they quickly reacted to. Since there were 16 rewarded trials and 4 oddball trials, the total bonus amounted to 4 £ and they received 1 £ for each oddball trial that they reacted to. In line with the appetitive reinforcement we collected reward expectation ratings instead of shock expectations. In each rating subjects were presented with every face for 1.5 seconds. Afterwards they were shown a 10-point Likert scale with the same anchors as in studies 1 and 2. The question with respect to which subjects were instructed to respond was „How likely is this face going to reward you with money?”. Subjects chose their response with the left mouse key and submitted their response by clicking the “continue” button. We used the same sequences and ITIs that we also used in the previous studies.

### Model fitting

All behavioral models were implemented as hierarchical Bayesian models in the Stan probabilistic programming language (38) using `pystan`. We used priors that we considered weakly informative in all models. The specific priors are described with the models. To avoid divergent transitions, we used a non-centered parameterization of all hierarchical parameters (39). All models were run for 4 chains until those converged. Typically this meant 2000 iterations each, where the first 1000 samples per chain were treated as warmup and discarded. We increased the number of samples in steps of 1000 as needed. Convergence was defined as all 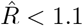 [82]. For transparency, all model fits are included in the data.

### Bayesian model comparison

To compare different Bayesian models we used approximate leave-one-out cross-validation (LOO-CV), which is a method to compute out of sample predictive accuracy via the expected log predictive density (ELPD) of left-out data points. The specific version we employed [83] requires sampling from an auxiliary mixture distribution, which in contrast to purely post-processing based approaches [84] is robust to very informative data points. Note that this method does not require explicit penalization because adding parameters in a Bayesian model distributes the probability over more dimensions and results in lower pointwise probability density in the prior, which acts as a built-in penalty. In addition, LOO-CV avoids overfitting by using out of sample predictions. We considered a model to fit the data credibly better than another model if the ELPD was higher and the difference of the ELPDs was larger than its standard error.

### Hierarchical mean posterior difference scaling (hMPDS)

To fit the perceptual spaces we adapted MLDS [80], which is a method to fit one-dimensional psychometric functions. In earlier work from our lab [16] it has been generalized to multiple dimensions and was used to fit perceptual spaces. In MLDS, positions of stimuli ***Ψ*** and an estimator for the standard deviation of perceptual noise *σ* are optimized to maximize the likelihood of subjects’ decisions in a quadruplet task (see above). The quantities being compared in the likelihood function are the within-pair perceptual distances of both pairs given the positions ***Ψ***, i.e. the length of the difference vector:

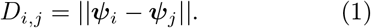

For the likelihood function, the difference between the two distances of the two pairs is normalized using *σ* and - assuming Gaussian perceptual noise - entered into the standard normal CDF to give the likelihood of choosing the second pair (coded as 1):

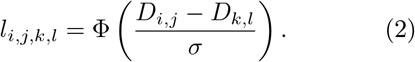

The likelihood function of a single observation is then just the Bernoulli PMF

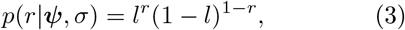

where we omit the subscripts for the stimuli for readability. The likelihood of a vector of independent observations ***r*** is the product of the likelihoods of the individual observations. We implemented a hierarchical Bayesian version, which we call hMPDS, to exploit the fact that perceptual spaces are unlikely to be completely independent, i.e. the perception of different people is somewhat similar, and to include prior knowledge from the construction of the stimulus set. The likelihood function of hMPDS is exactly the MLDS likelihood function. The subject level stimulus positions *ψ*_*s*_ are assumed to be normally distributed around the group level positions *ψ*_*g*_ with standard deviation *σ*_*Ψ*_ for each stimulus i, dimension d and subject s:

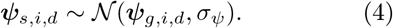

We used weakly informative priors on the group level stimulus positions with the mean being chosen according to an optimal 3 × 3 grid with positions going from 0 to 1. I.e. the lower left stimulus had group level priors

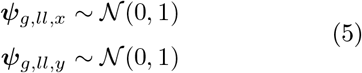

for the x and y position while the priors for the upper left stimulus were

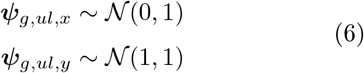

Without constraints, this model is only partially identified through the priors as the solution is invariant to translation, scaling, rotation and reflection. Intuitively, because the likelihood only depends on the within-pair distances, the likelihood of an estimate for *Ψ* is the same if one shifts, reflects or rotates the solution as these distances stay the same. In order to fully identify the model, we fixed two stimulus positions and constrained one x-position to be on the left of the line defined by the two fixed positions while leaving all subject level positions unconstrained. The perceptual noise was sampled on the log scale and subject level log noise terms log(*σ*_*s*_) were assumed to be normally distributed around the group level log noise log(*σ*_*g*_) with standard deviation *σ*_*σ*_:

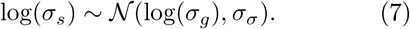

To control for changes in the perceptual space of subjects we compared two different models in studies 1 and 2. In the first version (Dynamic), perceptual spaces before and after the experiment were independent sets of parameters, i.e. the model allowed for changes in perceptual spaces due to the generalization experiment. In the second version (Fixed), the spaces were constrained to be the same, assuming no change to the spaces due to conditioning. In both studies and conditions the constrained version fits the data better than the version with separate perceptual spaces (tab. S1). Thus, we conclude that perceptual spaces stay approximately constant and asymmetric generalization is not due to distortions in this space that were introduced by the learning experience. Posterior means were used as estimators for the stimulus positions in perceptual space. To be able to include the perceptual spaces in the analysis, we interpolated the stimulus positions using a second-order multivariate polynomial regression as implemented in the python package `scikit-learn`. In particular, we computed the mapping from the polynomial features of the optimal 3 × 3 grid to the fitted posterior means from hMPDS on an individual level. This mapping was applied to the complete optimal 5 × 5 grid. The solution was procrustes aligned to the optimal 5 × 5 grid to generate the full individual perceptual space.

### Bayesian model of generalization

In our model, agents learn an exponential mapping from psychological space onto outcome probabilities, which we refer to as associative maps in analogy to Lee *et al*. [47]. In *n* dimensions, this mapping is characterized by the *n*-dimensional vectors ***μ*** and ***λ***, which are the midpoint and the strength of exponential decay respectively and the scalar *ρ*, which is the outcome probability at ***μ***. Any set of these parameters defines an associative map. The choice of an exponential shape is based on Shepard’s [1] observations, but it is a relatively arbitrary one and in principle the model could be extended to other shapes or even a probability distribution over possible shapes. The likelihood of observing a consequential outcome given this mapping depends on the weighted cityblock distance *δ* along the n dimensions. For a stimulus at position *s*_*i*_ this is given by

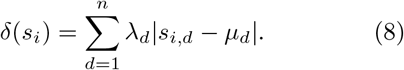

Note that changing the decay parameter *λ* for a dimension has the exact same effect as rescaling the respective dimension which directly includes partial dimensionality reduction and full reduction as *λ* approaches 0. For integral stimulus dimensions, this could be replaced with a weighted euclidean distance that relies on a single parameter as a preference for one of several dimensions that cannot be separated seems impossible.

Because perceptual noise is an important consideration with respect to the debate about its role in generalization in cognitive neuroscience and because it acts as a lower bound on the width of generalization, we constructed a symmetric k x k perceptual confusion matrix ***A***, where *k* = 25 is the number of stimuli. The probability of confusing stimuli i and j was given by a multivariate normal distribution with covariance matrix **Σ** = *ϵ*^2^**I**, where *ϵ* is the standard deviation of Gaussian perceptual noise and **I** is the identity matrix.

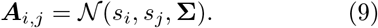

The rows of ***A*** were normalized to sum to 1 in order to account for the discretization of the continuous stimulus space. Multiplying the vector ***δ*** of distances with the perceptual confusion matrix ***A*** yields an estimate of the perceptual distances between stimuli and ***μ*** that accounts for perceptual noise:

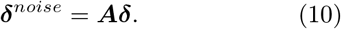

Finally, the probability of observing a consequence r given the parameters ***θ*** = {***μ, λ***, *ρ*} and after observing a stimulus s decays with perceptual distance to ***μ*** and is equal to *ρ* at the noise-free highest point:

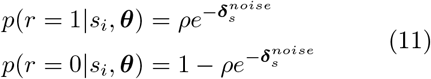

which is the likelihood that is used to update the belief state about model parameters. Due to the noisy estimate of distance to the center of the association map, the inferred probability for a stimulus is always lower than *ρ*, which corresponds to a lapse rate in other models. Note that this likelihood function corresponds to weak sampling [28]. Factoring the likelihood into *p*(*s, r*|***θ***) = *p*(*r*|*s*, ***θ***)*p*(*s*|***θ***) allows for an inclusion of other sampling assumptions by substituting an appropriate equation for *p*(*s*|***θ***). But since weak sampling seems like the best-fitting assumption in our experiments as we randomly sample from the stimulus space, the term is constant and drops from the equation, which results in eq. 11 for the likelihood.

### Generating predictions

For our simulations, we assumed our stimuli to be arranged in a 5 × 5 grid with a range of (0, 1) for the emotion dimension and (−0.5, 0.5) for the identity dimension. The full space we considered had a range of (0, 1) on the emotion dimension and (−1, 1) on the identity dimension. The reason for this choice is that while the emotion dimension has natural endpoints (a completely neutral and a maximally angry or happy face), the identity dimension does not. Faces that are more dissimilar from each other are imaginable. This is an important consideration because a uniform prior on the center of the consequential region or association map does not imply a uniform generalization gradient if the considered space is constrained [30]. In the case of consequential region models this is because stimuli in the center of the considered space are part of more consequential regions. In our case, stimuli in the center are on average closer to the midpoint of associative maps which predicts a higher a priori outcome probability for those stimuli. On the emotion dimension this effect is counteracted by the informed priors, on the identity dimension it is attenuated by considering values for *μ*_*identity*_ that are outside of the range of stimuli. For the simulation we assumed a uniform prior on the midpoint parameters of the identity dimension:

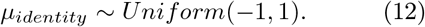

On the emotion dimension we used an informative Beta prior. To account for the intuition that angry faces are more likely to predict an aversive and less likely to predict an appetitive outcome while this pattern is flipped for happy faces, we put strongly informative priors on *μ*_*emotion*_, the midpoint parameters of the emotion dimension. For the context of aversive conditioning we used

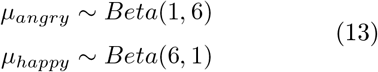

and in the context of appetitive conditioning, we flipped the parameters:

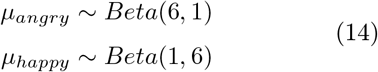

We also used a Beta prior on the shock probability that is skewed towards smaller outcome probabilities to account for the instructions that read that reinforcement would happen “occasionally”.

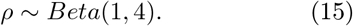

Dimensionality reduction or a dimension preference as predicted by representation learning corresponds to different priors on ***λ*** for both dimensions in the context of our model. To account for the prior knowledge on the relevance of dimensions we used

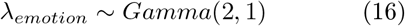

on the emotion dimension and

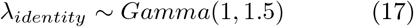

on the identity dimension, where Gamma is parameterized following Stan convention using shape and inverse scale. This choice shifts the prior towards smaller values and increases certainty on the identity dimension. Intuitively this means that the identity dimension is not considered to be very relevant with respect to the outcome and an increased amount of counterfactual evidence is needed to overwrite this belief state. Importantly, a decrease in the decay parameter has the exact same effect as a downscaling of that dimension, which naturally includes dimensionality reduction. Finally, the most extreme version of the aforementioned effect would be a complete neglect of facial identity which would result in full dimensionality reduction to a one-dimensional psychological space. We include this possibility because it would suggest an account of dimensionality reduction that is not compatible with our Bayesian model. We implemented this as a model in which *μ* and *λ* were scalars instead of vectors and the only position of stimuli considered was the position along the emotion axis.

### Bayesian model predictions

A value of *ϵ* = 0.05 was used for the perceptual noise variance for all simulations because it suggests a small probability of confusing neighboring faces, which matched our intuition. To generate predictions for all five ratings, we implemented this model in Stan and trained it on the same sequence of shocks or rewards and stimuli that we presented human subjects with. For the first rating we included no data, i.e. we sampled from the prior. For the second rating we included the sequences of stimuli and shocks of the first mesoblock, for the third rating sequences for the first two mesoblocks and so on. Note that in theory it would be possible to use the posterior after observing the first mesoblock as a prior for the next mesoblock. However, in the absence of a closed form solution, we opted to retrain the model with all the stimuli and shocks that were observed up to the rating at hand, which is equivalent. Generating predictions from this model for shock or reward expectation ratings relies on the posterior distribution on model parameters

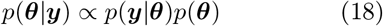

and the probability of a new observation given the model parameters and previous observations *p*(*ŷ*|***y, θ***), where ***y*** = {***s, r***} is the set of all observations and 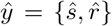 is the new observation. Predictions for future observations are generated by integrating the probability of new data over the posterior:

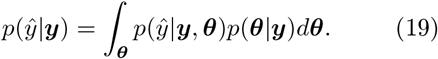

Formally, eq. 19 refers to the probability of observing a pair of a stimulus and outcome. But since ratings are collected with respect to a specific stimulus, i.e. after observing the stimulus, this is equivalent to the probability of an outcome given the stimulus. Note that the joint probability of outcome and stimulus can be factored into

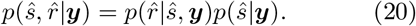

and since *p*(*ŝ*|***y***) = 1 after observing it, this simplifies to

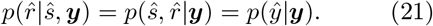

Since we don’t have a closed-form solution for the posterior, we rely on MCMC to sample from it and approximate expectations by averaging over samples from the posterior *p*(***θ***|***y***):

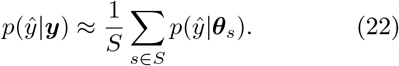

The prior distribution *p*(***θ***) corresponds to the prior belief state of subjects and itself depends on hyperpriors which we fixed in the simulations. These are the parameters of the priors on ***μ, λ*** and *ρ*. If we were to fit the model to data, we would try to computationally characterize the prior assumptions of subjects, i.e. those hyperpriors would be the parameters we would be interested in. Thus, the likelihood function we would try to optimize or use in a Bayesian model is the likelihood of shock or reward expectation ratings ***z*** given the hyperpriors ***τ***. Treating hyperpriors as free parameters induces a new dependency on ***τ*** in eq. 19 and the predicted probability of future events given the hyperpriors is given by

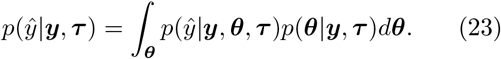

which assuming the conditional independence

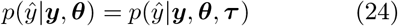

simplifies to

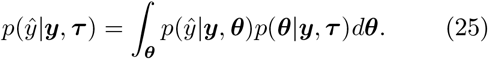

And again, since we are using a sampling algorithm, we would have to approximate this quantity like so:

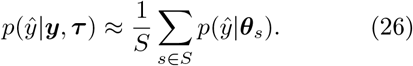

with ***θ***_*s*_ being sampled according to the conditional posterior *p*(***θ***|***τ***, *y*). In addition, ratings were given on a scale from 1-10 and include measurement errors and response tendencies. This means that we would need an error model for the relationship between posterior expectations and ratings in order to derive a likelihood function. As a consequence, we would need a mapping from *p*(*ŷ*|*y*, ***τ***) to observed shock or reward expectation ratings, e.g. a linear mapping

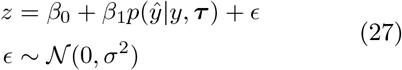

which would yield the likelihood

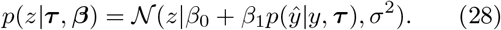

Note that since the model is already written in Stan, we cannot use the same language to fit it to data because we would be sampling from a joint posterior *p*(*z, y*|***θ, τ***) that depends on both the likelihood of ratings given the implied outcome probabilities and the likelihood of stimuli and shocks given the prior assumptions. The practical problem of fitting this model lies in the fact that *p*(*ŷ*|***y, τ***) is very expensive to compute and subject to approximation errors due to relying on sampling which needs to be recomputed for every iteration, which yields a very hard optimization problem. Since our attempts to still fit the model using Bayesian Optimization (BO) with different surrogate models did not yield a converging solution within a reasonable amount of time, we did not fit the full Bayesian model to the data. Importantly, this is not required for the model to be useful and we followed previous examples in the literature by using it to make qualitative predictions about empirical data and derive specific hypotheses that we test using a simpler model that can be fit to data [31, 47, 85]. This way we build a bridge between cognitive models in rational analysis, representation learning and more empirically focussed work in stimulus generalization while having a theory and a generative mechanism in mind when interpreting simpler fitted models instead of purely describing the data. These models are described in detail below.

### Fitted models

We heuristically derived three different approximations to the full Bayesian model that allowed us to test the different hypotheses using posterior parameter estimates and model comparison. The three models differ only in the notion of dimensionality reduction they include. We compared models that assume either no („No DR”), full („Full DR”) or partial („Partial DR”) dimensionality reduction. While fitting these models, we included hierarchically fitted single subject perceptual spaces to determine emotionality (abbreviated as *emo*), identity (*id*) and proximity either as Euclidean distance or separately along both dimensions (*pemo* and *pid*) to the reinforced stimulus for all stimuli. This was done in order to control for perceptual differences between neighboring faces across both dimensions and to prevent a putative effect of distortions in perceptual spaces. All parameters of these models were implemented hierarchically assuming normal distributions of individual parameters around the group level parameters and weakly informative priors on the group level. The specific priors we used are reported below. Posterior predictive checks on the group level were computed by averaging model-predicted ratings over samples and subjects to ensure that all subjects contribute equally to the expectation as they did for the empirical mean ratings. The same would not be true when computing the ratings that would be implied by the group level parameters because the influence of subjects on the group level parameters depends on the precision of the estimation of single subject parameters.

### No DR

The first hypothesis from the Bayesian model is that ratings are initially driven by the amount of emotional expression in the stimuli. With ongoing learning the proximity to the reinforced (center) stimulus should become more important while the importance of the emotional expression should decrease. Another unlikely possibility would have been that one of the identities that we morphed into each other to create an identity continuum could have been considered more dangerous a priori. This model excludes the possibility of dimensionality reduction because the measure of proximity to the CS+ is the Euclidean distance in perceptual space. To formalize this idea, we fit a model that predicted the different ratings at every time point as a combination of a baseline shock expectation and the influences of emotionality, identity and proximity to the reinforced stimulus: The predicted rating 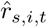 for subject s, stimulus i at time t was modeled as

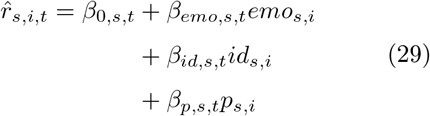

The subscript for subjects on emo, id and p is needed because we included individual perceptual spaces. To account for the predictions that the relative influence of these features changes with time and that the updating from rating to rating should decrease, we modeled time dependence with an intercept and slope per feature and added another term *λ* that allowed for non-linear changes in the influence of the features over time. In particular, the influence of the features at time *t* ∈ {0, 1, 2, 3, 4} was modeled as

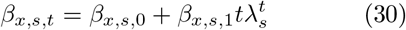

where *x* is a placeholder for the different features. This corresponds to an increase in updating for *λ >* 1 and a decrease for *λ <* 1. Note that this model can actually predict a return to baseline and below for small values of *λ*, but we did not expect this to be a large issue for the present application.

Lastly, we hypothesized that with increasing certainty in the belief state (i.e. certainty that only a small region in psychological space predicts a shock), the estimated exponential decay in the associative map in the Bayesian model gets stronger. To include this in the model we added an exponential term *ρ* that is itself time-dependent and governs the slope of perceptual distance:

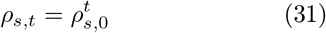

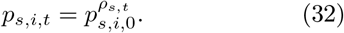

Here, *p*_*s,i*,0_ is the perceptual distance as implied by the fitted perceptual. This parameter implies increasingly wider generalization around the reinforced stimulus for *ρ <* 1 and increasingly narrower generalization for *ρ >* 1. Ratings were assumed to be normally distributed around the predicted rating with variance *σ*_*rating*_.

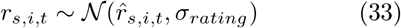

**Priors** As group-level priors, we used

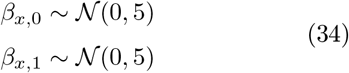

on the intercept and slope of the time-dependent influence of the features and on the slope of the baseline shock expectation and

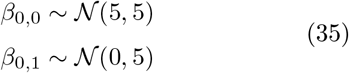

on the intercept of the baseline shock expectation. Because even small deviations from 1 have a strong effect for the non-linear parameters *λ* and *ρ*, we used narrower priors on those that still allowed for large deviations from 1:

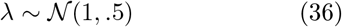

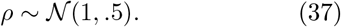

The standard deviation of ratings around predicted ratings was fit per subject without hierarchical dependence and sampled on the log scale with a

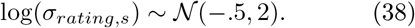

prior. As priors on hierarchical standard deviation parameters we used

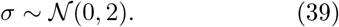

for all *β*s and

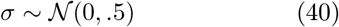

for *λ* and *ρ*. Note that these are half-normal distributions because those parameters are by definition positive.

### Partial DR

To include the idea of partial dimensionality reduction, that is a preference for one over the other dimension due to a rescaling of perceptual spaces, we fit a second model in which proximity to the CS+ is split into two dimensions. In particular, for this model eq. 29 was replaced by

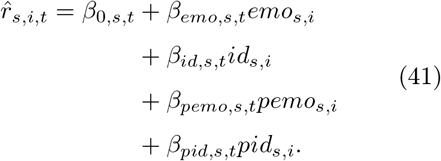

Analogously, eq. 32 was replaced by

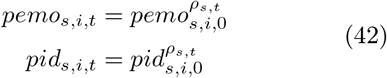

### Full DR

Lastly, we investigated the possibility of a complete neglect of the identity dimension. For this reason, we included a model that only contains information about the emotional dimension. In this model, the predicted rating and the time dependent perceptual distance are computed as

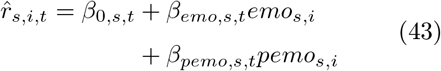

and

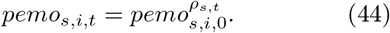

### fMRI acquisition

Functional and anatomical MR images were collected using a 3T PRISMA MRI Scanner (Siemens, Erlangen, Germany) with a 64 channel head coil. For the functional scans a multiband EPI sequence was used (multiband factor = 3, TR = 1.526s, TE = 30ms, flip angle = 60°, FOV = 225mm, GRAPPA PAT-factor: 2, reference lines: 48). Each volume consisted of 63 slices with an isotropic voxel size of 1.5mm and no gap between slices. In order to account for B0 field inhomogeneity a B0 field map was collected before each MRI session (63 slices, TR = 772, short TE = 6.21 ms, long TE = 8.67, FOV = 222mm, flip angle = 40°). In addition, for each subject a high-resolution (voxel size = 1mm isotropic) anatomical T1-weighted image was acquired using an MPRAGE sequence.

### fMRI preprocessing

All MRI data was analyzed using `SPM12` (Wellcome Trust Center for Neuroimaging, London, UK) and `Matlab` (Mathworks). Before preprocessing, the first four scans per run were discarded. Preprocessing consisted of motion correction (realignment and field map correction) and slice time correction. Mean EPI images per subject were calculated and segmented to generate native tissue class images. These tissue class images were used to create a flow field of the mean EPI in MNI standard space (IXI555_MNI152 template of the `CAT12` toolbox) using the DARTEL toolbox in SPM12. We did not normalize functional images to MNI space, but instead computed first level analysis in the native space and then normalized and smoothed the resulting beta images with a Gaussian kernel (FWHM=6mm).

### Univariate fMRI analysis

Univariate analysis of fMRI data consisted of two GLMs. For both GLMs we defined onset regressors at the onsets of the stimulus presentation and the electrical stimulation respectively. We combined both data sets for the fMRI analysis to increase statistical power and identify mechanisms that are agnostic to the emotional expression used. That is, we were interested in the neural correlates of the impact of prior knowledge as a general concept, not constrained to the case of either emotion. All first level analysis included functional scans from four experimental runs (see design). Besides the regressors of interest we entered one intercept per run and six movement regressors into the GLMs. Beta images of the first level analysis were warped into MNI space using the flow fields generated from the individual tissue class images and smoothed before entering them into the second level analysis. For the model-based fMRI analysis we used predictions from the fitted model.

### GLM1

For the first GLM we interpolated the fitted ratings on an individual level to generate approximate model-implied shock expectations for the single trials. In particular, in the behavioral model, the five ratings blocks were modeled using the time vector [0, 1, 2, 3, 4]. To generate predictions for the 20 microblocks of the experiment, we computed posterior predictive distributions for ratings given a time vector consisting of 20 equidistant values from 0 to 4. Note that this approach assumes that the belief state about the predictive value of stimuli is constant within microblocks due to constraints in the modeling approach. While this is technically incorrect, it represents a good approximation. Posterior means of interpolated shock expectations were placed as parametric modulators on the onset of stimuli. Beta images from the first level analysis were entered into a second level fixed effect ANOVA as implemented in `SPM12`. After estimating the model we computed positive and negative t-contrasts to identify areas with a positive and negative generalization tuning respectively. We applied a whole brain family wise error rate (FWE) with a threshold of *α* = .025, which corresponds to a two-sided t-test at *α* = .05. Two-sided T-images for visualization were computed as the difference between the thresholded T-maps of the positive and negative contrast. The threshold for the uncorrected t-contrasts for visualization was set to .0005, corresponding to a two-sided *α*-level of .001.

### GLM2

The approach for the second GLM was analogous to the previous approach, except that we interpolated the features that make up the overall shock expectation individually instead of the full fitted ratings. These features comprise a baseline shock expectation term (*β*_0,*s,t*_), the impact of emotionality (*β*_*emo,s,t*_*emo*_*s*_) and identity (*β*_*id,s,t*_*id*_*s*_) of faces and the impact of the perceptual proximity along the emotion (*β*_*pemo,s,t*_*pemo*_*s*_) and identity (*β*_*pid,s,t*_*pid*_*s*_) dimensions. All features were entered as parametric modulators on the stimulus onset regressors. For the second level analysis we entered all beta images into another ANOVA and computed positive and negative t-contrasts for each parametric modulator. We opted to use directed contrasts for all features because for the parametric modulators that indicate the impact of perceptual proximity to the reinforced stimulus we wanted to identify areas that show a positive or negative perceptual tuning and for all other modulators we did not have directed hypotheses but were interested in the direction of the effect. Two-sided T-maps were computed as described above and the *α*-level was set to 0.25 to account for two-sided testing.

### Representational Similarity Analysis

For the representational similarity analysis we computed single-trial beta images using a least squares separate approach [86]. This approach requires fitting one GLM per trial in which there is one regressor for the trial of interest and one regressor in which all other trials are collapsed. We constructed model RDMs using the individually fitted perceptual spaces of subjects. For the Emotion RDM, we defined the dissimilarity between stimuli as the absolute distance between the position estimates along the emotion axis for each subject. For the Identity RDM, we did the same thing along the identity axis. For the searchlight analyses, we used the `brainiak Python` package. For each voxel the searchlight considered all voxels that were within a radius of 5mm. Neural dissimilarities were computed using correlation distance. For the similarity between neural and model RDMs we used spearman correlations. First, we ran whole brain searchlight analysis using beta images that were averaged over the whole experiment per stimulus (20 beta images per stimulus) for the emotion and identity RDM separately. For group level inference, spearman correlations were Fisher-Z transformed. Those Z-maps were normalized to MNI space, smoothed using a 6mm Gaussian Kernel and entered into a secondlevel fixed effect ANOVA using `SPM12`. For the secondlevel model, we computed a directed T-contrast to look for areas that had higher correlations with the emotion RDM then the identity RDM.

We repeated the analysis using beta images that were averaged over stimuli within run, resulting in 4 whole brain searchlights per model RDM. All Fisher-Z images were treated as above and jointly entered in a secondlevel fixed effect ANOVA. We computed an F-contrast with one row per block and condition (emotion vs. identity) while averaging over sessions (angry vs. happy) to look for brain areas that showed significant correlations with either the emotion or identity RDM in at least one run, independently of the emotion. We then extracted single subject Fisher-Z transformed correlations from the peak voxels of this analysis in bilateral MFG and left IPS and further analyzed these with Bayesian hierarchical linear regressions to characterize the time course. We chose linear regressions despite the fact that correlations are bounded between −1 and 1 because of the simplicity, the ease of interpretation of parameter estimates and because the hard constraints on values were unlikely to play a role within the range of values that we observed. The regression included two levels of hierarchy: conditions (angry vs. happy) in subject and subjects in group. The dependent variable were the Fisher-Z transformed spearman correlations *ρ* between the neural RDMs, centered on the peak voxel and the model RDMs (emotion or identity). Independent variables were time points t, ranging from 0 for the first run to 3 for the last run and a vector of ones for the intercept. That is, the equation we fitted was

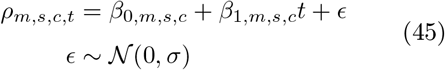

where s indicated subject, c indicates condition (angry vs. happy) and m indicated the model RDM (emotion vs. identity). Subject-level parameters were normally distributed around group level parameters

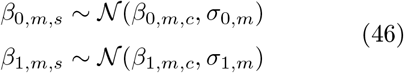

Individual condition level parameters were normally distributed around subject level estimates with a common standard deviation

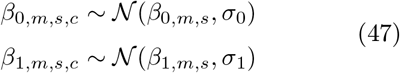

Due to the noisy nature of the dependent variables and the observed small correlations, we used narrow *𝒩* (0, 0.1) priors on all group level parameters and hierarchical standard deviations and a standard normal prior on *σ*. Note that the comparison of prior and posterior standard deviations indicated that those priors were not overly informative since

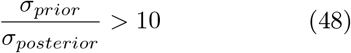

for all group level parameters.

### Brain behavior correlation

For the correlation between the regression coefficients of the RSA correlations and the behavioral model parameters we computed the posterior mean over those parameters. In particular, we extracted *β*_1,*id,s,c*_ for all subjects and both conditions from the hierarchical regression of RSA correlations and *β*_*pid,s*,1_ from the fitted behavioral models in both conditions and computed Pearson correlations separately for both conditions.

### Software

Experimental scripts were programmed in `Matlab` and `psychtoolbox3` for studies 1 and 2 and in `HTML` and `Javascript` using the `jspsych` package for study 3. fMRI data was analyzed in `Matlab` and `SPM12` except for the RSA search-light analyses, for which we used the `brainiak` package in `Python`. For the least squares separate GLM that we used to create beta images for the RSA analysis, we used code by Selim Onat (retrieved from https://github.com/selimonat/fancycarp/blob/mrt/fearamy/). Bayesian models were written in `Stan` which we interfaced from `Python` using the `pystan` package. Everything else was done using `Python` including the third-party packages `numpy`, `scipy`, `pandas`, `matplotlib`, `seaborn`, `nilearn`, `nibabel`, `scikit`-learn, `arviz` and `numba`.

## Acknowledgements

We would like to thank Michael Frank and Nathaniel Daw for helpful comments in the planning state of the presented work, Estefanía Orozco Rodríguez and Jenny Luca Rohwer for their assistance in data collection, Selim Onat for inspiring the study design and for code for the least squares separate fMRI analysis and Marie Habermann and Lea Kampermann for helpful input on an earlier version of the manuscript.

## Funding

This work was funded by the Landesforschungs-förderung Hamburg and ERC AdG-883892-PainPersist.

## Data and code availability

All code and data is publicly available at https://gin.g-node.org/LukasNeugebauer/neugebauer-buechel-2023/src/main.

## Author contributions

LMN: Conceptualization, Methodology, Software, Data analysis, Investigation, Visualization, Project administration, Writing – original draft, Writing – review & editing. CB: Conceptualization, Methodology, Funding acquisition, Project administration, Supervision, Writing - review & editing

## Competing interests

The authors declare that they have no competing interests.

## Notes

### Competing Interest Statement

The authors have declared no competing interest.

## References

1. Shepard, R. N. Toward a universal law of generalization for psychological science. Science 237, 1317–1323.

2. Webler, R. D. et al. The neurobiology of human fear generalization: meta-analysis and working neural model. Neuroscience & Biobehavioral Reviews 128, 421–436.

3. Dymond, S., Dunsmoor, J. E., Vervliet, B., Roche, B. & Hermans, D. Fear Generalization in Humans: Systematic Review and Implications for Anxiety Disorder Research. Behavior Therapy 46, 561–582.

4. Soto, F. A. & Wasserman, E. A. Integrality/Separability of Stimulus Dimensions and Multidimensional Generalization in Pigeons. Journal of experimental psychology. Animal behavior processes 36, 194–205.

5. Guttman, N. & Kalish, H. I. Discriminability and stimulus generalization. Journal of Experimental Psychology 51, 79–88.

6. Blough, D. S. Steady state data and a quantitative model of operant generalization and discrimination. Journal of Experimental Psychology: Animal Behavior Processes 1, 3–21.

7. Ghirlanda, S. & Enquist, M. A century of generalization. Animal Behaviour 66, 15–36.

8. Gerraty, R. T., Davidow, J. Y., Wimmer, G. E., Kahn, I. & Shohamy, D. Transfer of Learning Relates to Intrinsic Connectivity between Hippocampus, Ventromedial Prefrontal Cortex, and Large-Scale Networks. Journal of Neuroscience 34, 11297–11303.

9. Niv, Y. Learning task-state representations. Nature Neuroscience 22, 1544–1553.

10. Voorspoels, W., Navarro, D. J., Perfors, A., Ransom, K. & Storms, G. How do people learn from negative evidence? Non-monotonic generalizations and sampling assumptions in inductive reasoning. Cognitive Psychology 81, 1–25.

11. Yantis, S. The Neural Basis of Selective Attention: Cortical Sources and Targets of Attentional Modulation. Current Directions in Psychological Science 17, 86–90.

12. Jackson, J., Rich, A. N., Williams, M. A. & Woolgar, A. Feature-selective Attention in Frontoparietal Cortex: Multivoxel Codes Adjust to Prioritize Task-relevant Information. Journal of Cognitive Neuroscience 29, 310–321.

13. Fusi, S., Miller, E. K. & Rigotti, M. Why neurons mix: high dimensionality for higher cognition. Current Opinion in Neurobiology. Neurobiology of cognitive behavior 37, 66–74.

14. Bottini, R. & Doeller, C. F. Knowledge Across Reference Frames: Cognitive Maps and Image Spaces. Trends in Cognitive Sciences 24, 606–619.

15. Spence, K. W. The differential response in animals to stimuli varying within a single dimension. Psychological Review 44, 430–444.

16. Onat, S. & Büchel, C. The neuronal basis of fear generalization in humans. Nature Neuroscience 18, 1811–1818.

17. Schechtman, E., Laufer, O. & Paz, R. Negative Valence Widens Generalization of Learning. Journal of Neuroscience 30, 10460–10464.

18. Laufer, O., Israeli, D. & Paz, R. Behavioral and Neural Mechanisms of Overgeneralization in Anxiety. Current Biology 26, 713–722.

19. Laufer, O. & Paz, R. Monetary Loss Alters Perceptual Thresholds and Compromises Future Decisions via Amygdala and Prefrontal Networks. Journal of Neuroscience 32, 6304–6311.

20. Zaman, J., Struyf, D., Ceulemans, E., Beckers, T. & Vervliet, B. Probing the role of perception in fear generalization. Scientific Reports 9, 10026.

21. Struyf, D., Zaman, J., Hermans, D. & Vervliet, B. Gradients of fear: How perception influences fear generalization. Behaviour Research and Therapy 93, 116–122.

22. Struyf, D., Zaman, J., Vervliet, B. & Van Diest, I. Perceptual discrimination in fear generalization: Mechanistic and clinical implications. Neuroscience and Biobehavioral Reviews 59, 201–207.

23. Greenberg, T., Carlson, J. M., Cha, J., Hajcak, G. & Mujica-Parodi, L. R. Ventromedial Prefrontal Cortex Reactivity Is Altered In Generalized Anxiety Disorder During Fear Generalization. Depression and Anxiety 30, 242–250.

24. Lissek, S. et al. Overgeneralization of Conditioned Fear as a Pathogenic Marker of Panic Disorder. The American journal of psychiatry 167, 47–55.

25. Lissek, S. Toward an Account of Clinical Anxiety Predicated on Basic, Neurally Mapped Mechanisms of Pavlovian Fear-Learning: The Case for Conditioned Overgeneralization. Depression and Anxiety 29, 257–263.

26. Kaczkurkin, A. N. et al. Neural Substrates of Overgeneralized Conditioned Fear in PTSD. American Journal of Psychiatry 174, 125–134.

27. Anderson, J. R. The adaptive character of thought ISBN: 978-0-8058-0419-5 (Lawrence Erlbaum Associates, Inc, Hillsdale, NJ, US).

28. Tenenbaum, J. B. & Griffiths, T. L. Generalization, similarity, and Bayesian inference. Behavioral and Brain Sciences 24, 629–640.

29. Navarro, D. J., Dry, M. J. & Lee, M. D. Sampling Assumptions in Inductive Generalization. Cognitive Science 36, 187–223.

30. Navarro, D. J., Lee, M. D., Dry, M. J. & Schultz, B. Extending and Testing the Bayesian Theory of Generalization. Proceedings of the 30th Annual Conference of the Cognitive Science Society, 1746–1751.

31. Soto, F. A., Gershman, S. J. & Niv, Y. Explaining compound generalization in associative and causal learning through rational principles of dimensional generalization. Psychological Review 121, 526–558.

32. Sutton, R. S. & Barto, A. G. Reinforcement learning: an introduction Second edition. ISBN: 978-0-262-03924-6 (The MIT Press, Cambridge, Massachusetts).

33. Dayan, P. & Niv, Y. Reinforcement learning: The Good, The Bad and The Ugly. Current Opinion in Neurobiology. Cognitive neuroscience 18, 185–196.

34. Schultz, W., Dayan, P. & Montague, P. R. A neural substrate of prediction and reward. Science 275, 1593–1599.

35. Wimmer, G. E., Daw, N. D. & Shohamy, D. Generalization of value in reinforcement learning by humans. European Journal of Neuroscience 35, 1092–1104.

36. Wu, C. M., Schulz, E., Speekenbrink, M., Nelson, J. D. & Meder, B. Generalization guides human exploration in vast decision spaces. Nature Human Behaviour 2, 915–924.

37. Niv, Y. et al. Reinforcement Learning in Multidimensional Environments Relies on Attention Mechanisms. Journal of Neuroscience 35, 8145–8157.

38. Leong, Y. C., Radulescu, A., Daniel, R., DeWoskin, V. & Niv, Y. Dynamic Interaction between Reinforcement Learning and Attention in Multidimensional Environments. Neuron 93, 451–463.

39. Pettine, W. W., Raman, D. V., Redish, A. D. & Murray, J. D. Human generalization of internal representations through prototype learning with goal-directed attention. Nature Human Behaviour 7, 442–463.

40. Tomov, M. S., Dorfman, H. M. & Gershman, S. J. Neural computations underlying causal structure learning. The Journal of Neuroscience 38, 7143–7157.

41. Vaidya, A. R., Jones, H. M., Castillo, J. & Badre, D. Neural representation of abstract task structure during generalization. eLife 10 (eds Liljeholm, M., Ivry, R. B., Ranganath, C. & Michelmann, S.) e63226.

42. Eichenbaum, A., Scimeca, J. M. & D’Esposito, M. Dissociable Neural Systems Support the Learning and Transfer of Hierarchical Control Structure. Journal of Neuroscience 40, 6624–6637.

43. Schuck, N. W., Cai, M. B., Wilson, R. C. & Niv, Y. Human Orbitofrontal Cortex Represents a Cognitive Map of State Space. Neuron 91, 1402–1412.

44. Summerfield, C., Luyckx, F. & Sheahan, H. Structure learning and the posterior parietal cortex. Progress in Neurobiology 184, 101717.

45. Badre, D., Bhandari, A., Keglovits, H. & Kikumoto, A. The dimensionality of neural representations for control. Current Opinion in Behavioral Sciences. Computational cognitive neuroscience 38, 20–28.

46. Bernardi, S. et al. The Geometry of Abstraction in the Hippocampus and Prefrontal Cortex. Cell 183, 954–967.e21.

47. Lee, J. C., Lovibond, P. F., Hayes, B. K. & Navarro, D. J. Negative evidence and inductive reasoning in generalization of associative learning. Journal of Experimental Psychology: General 148, 289–303.

48. Keltner, D. & Kring, A. M. Emotion, Social Function, and Psychopathology. Review of General Psychology 2, 320–342.

49. Seligman, M. E. On the generality of the laws of learning. Psychological Review 77, 406–418.

50. Adolphs, R., Damasio, H., Tranel, D., Cooper, G. & Damasio, A. R. A Role for Somatosensory Cortices in the Visual Recognition of Emotion as Revealed by Three-Dimensional Lesion Mapping. Journal of Neuroscience 20, 2683–2690.

51. Zeidan, F., Lobanov, O. V., Kraft, R. A. & Coghill, R. C. Brain Mechanisms Supporting Violated Expectations of Pain. Pain 156, 1772–1785.

52. Drevets, W. C. et al. Blood flow changes in human somatosensory cortex during anticipated stimulation. Nature 373, 249–252.

53. Gijsen, S., Grundei, M., Lange, R. T., Ostwald, D. & Blankenburg, F. Neural surprise in somatosensory Bayesian learning. PLOS Computational Biology 17, e1008068.

54. Yeo, B. T. T. et al. The organization of the human cerebral cortex estimated by intrinsic functional connectivity. Journal of Neurophysiology 106, 1125–1165.

55. Andreatta, M. & Pauli, P. Generalization of appetitive conditioned responses. Psychophysiology 56, e13397.

56. FeldmanHall, O. et al. Stimulus generalization as a mechanism for learning to trust. Proceedings of the National Academy of Sciences 115, E1690–E1697.

57. Kampermann, L., Tinnermann, A. & Büchel, C. Generalization of placebo pain relief. Pain 162, 1781–1789.

58. Dunsmoor, J. E., Mitroff, S. R. & LaBar, K. S. Generalization of conditioned fear along a dimension of increasing fear intensity. Learning & Memory 16, 460–469.

59. Ghirlanda, S. Intensity generalisation: physiology and modelling of a neglected topic Journal (Paginated).

60. Zaman, J., Struyf, D., Ceulemans, E., Vervliet, B. & Beckers, T. Perceptual errors are related to shifts in generalization of conditioned responding. Psychological Research 85, 1801–1813.

61. Lommen, M. J. J., Engelhard, I. M. & van den Hout, M. A. Neuroticism and avoidance of ambiguous stimuli: Better safe than sorry? Personality and Individual Differences 49, 1001–1006.

62. Norbury, A., Robbins, T. W. & Seymour, B. Value generalization in human avoidance learning. eLife 7 (ed Lee, D.) e34779.

63. Van Meurs, B., Wiggert, N., Wicker, I. & Lissek, S. Maladaptive behavioral consequences of conditioned fear-generalization: A pronounced, yet sparsely studied, feature of anxiety pathology. Behaviour Research and Therapy 57, 29–37.

64. Ahmed, O. & Lovibond, P. F. The impact of previously learned feature-relevance on generalisation of conditioned fear in humans. Journal of Behavior Therapy and Experimental Psychiatry 46, 59–65.

65. Ahmed, O. & Lovibond, P. F. The Impact of Instructions on Generalization of Conditioned Fear in Humans. Behavior Therapy. Special Issue: Fear Generalization 46, 597–603.

66. Vervliet, B., Kindt, M., Vansteenwegen, D. & Hermans, D. Fear generalization in humans: Impact of verbal instructions. Behaviour Research and Therapy 48, 38–43.

67. Austerweil, J. L., Sanborn, S. & Griffiths, T. L. Learning How to Generalize. Cognitive Science 43, e12777.

68. De Voogd, L. D. et al. The role of hippocampal spatial representations in contextualization and generalization of fear. NeuroImage 206, 116308.

69. Goulden, N. et al. The salience network is responsible for switching between the default mode network and the central executive network: Replication from DCM. NeuroImage 99, 180–190.

70. Tuominen, L. et al. The relationship of perceptual discrimination to neural mechanisms of fear generalization. NeuroImage 188, 445–455.

71. Berg, H. et al. Salience and central executive networks track overgeneralization of conditioned-fear in post-traumatic stress disorder. Psychological Medicine, 1–10.

72. Woolgar, A., Thompson, R., Bor, D. & Duncan, J. Multi-voxel coding of stimuli, rules, and responses in human frontoparietal cortex. NeuroImage. Multivariate Decoding and Brain Reading 56, 744–752.

73. Flesch, T., Juechems, K., Dumbalska, T., Saxe, A. & Summerfield, C. Orthogonal representations for robust context-dependent task performance in brains and neural networks. Neuron 110, 1258–1270.e11.

74. Badre, D., Kayser, A. S. & D’Esposito, M. Frontal Cortex and the Discovery of Abstract Action Rules. Neuron 66, 315–326.

75. Stroud, J. P. et al. Ignorance is bliss: effects of noise and metabolic cost on cortical task representations. bioRxiv:2023.07.11.548492.

76. Rigotti, M. et al. The importance of mixed selectivity in complex cognitive tasks. Nature 497, 585–590.

77. Basu, R. et al. The orbitofrontal cortex maps future navigational goals. Nature 599, 1–4.

78. Jones, J. L. et al. Orbitofrontal cortex supports behavior and learning using inferred but not cached values. Science (New York, N.Y.) 338, 953–956.

79. Onat, S. Towards a new understanding of fear generalization and its neural origin. peerj:27311v1.

80. Maloney, L. T. & Yang, J. N. Maximum likelihood difference scaling. Journal of Vision 3, 5–5.

81. Watson, A. B. & Pelli, D. G. Quest: A Bayesian adaptive psychometric method. Perception & Psychophysics 33, 113–120.

82. Gelman, A. et al. Bayesian data analysis Third edition. ISBN: 978-1-4398-4095-5 (CRC Press, Boca Raton).

83. Silva, L. & Zanella, G. Robust leaveone-out cross-validation for high-dimensional Bayesian models. arXiv:2209.09190.

84. Vehtari, A., Simpson, D., Gelman, A., Yao, Y. & Gabry, J. Pareto Smoothed Importance Sampling. arXiv:1507.02646.

85. Behrens, T. E. J., Woolrich, M. W., Walton, M. E. & Rushworth, M. F. S. Learning the value of information in an uncertain world. Nature Neuroscience 10, 1214–1221.

86. Mumford, J. A., Turner, B. O., Ashby, F. G. & Poldrack, R. A. Deconvolving BOLD activation in event-related designs for multivoxel pattern classification analyses. NeuroImage 59, 2636–2643.

