## Supplementary figures and tables for "Bridging stimulus generalization and representation learning via rational dimensionality reduction"

### Supplementary Figures and Tables
### for
### "Bridging stimulus generalization representation learning via rational dimensionality reduction"

Lukas Michael Neugebauer, Christian Büchel

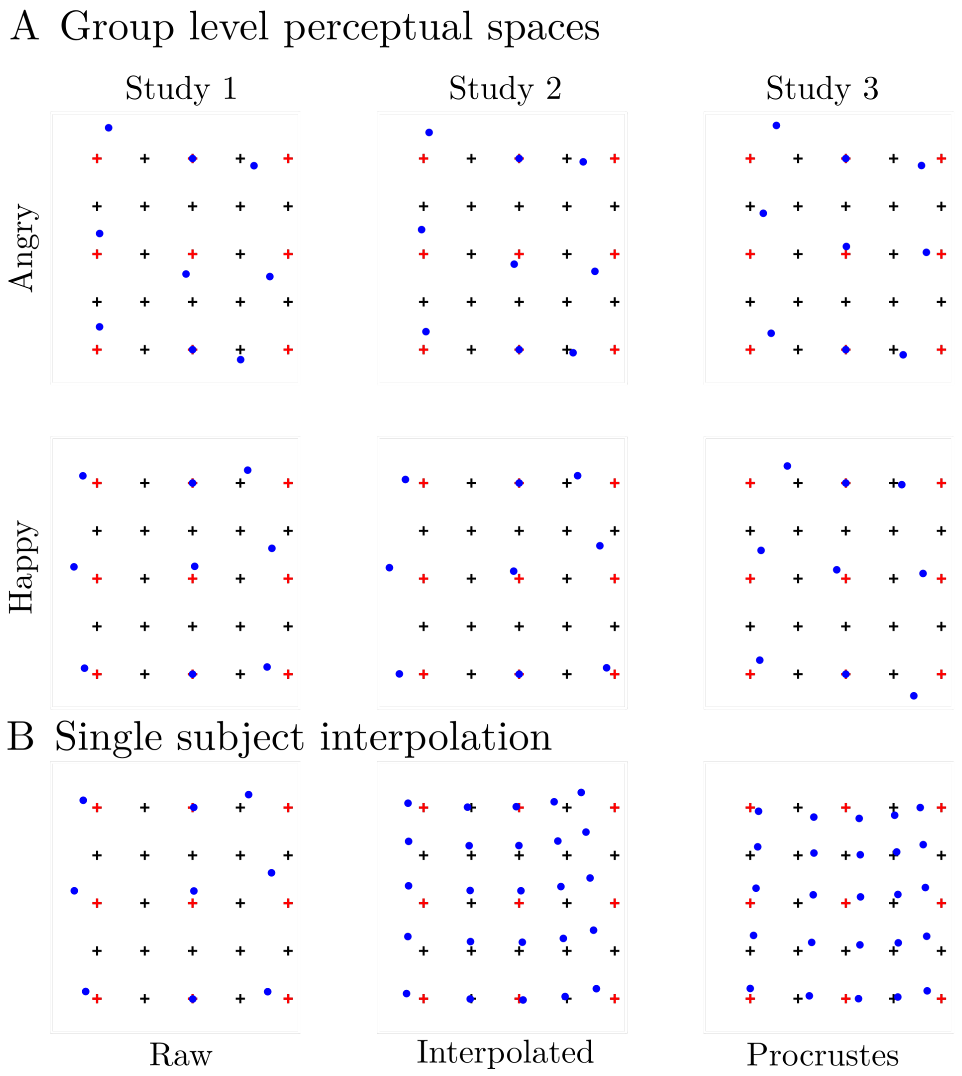


**Figure S1: Perceptual spaces.** (**A**) Group level fitted perceptual spaces were reasonable well aligned with the intended structure in all experiments (columns) and conditions (rows). (**B**) To include individual spaces into the analysis, we interpolated the fitted positions of 9 stimuli (column 1) to 25 stimuli (column 2) and procrustes aligned them to the optimal 5$x$5 grid (column 3). Note. Blue dots indicate the fitted positions, black crosses the optimal 5$x$5 grid and red crosses the optimal 3$x$3 grid of stimuli that were used for fitting the perceptual spaces.

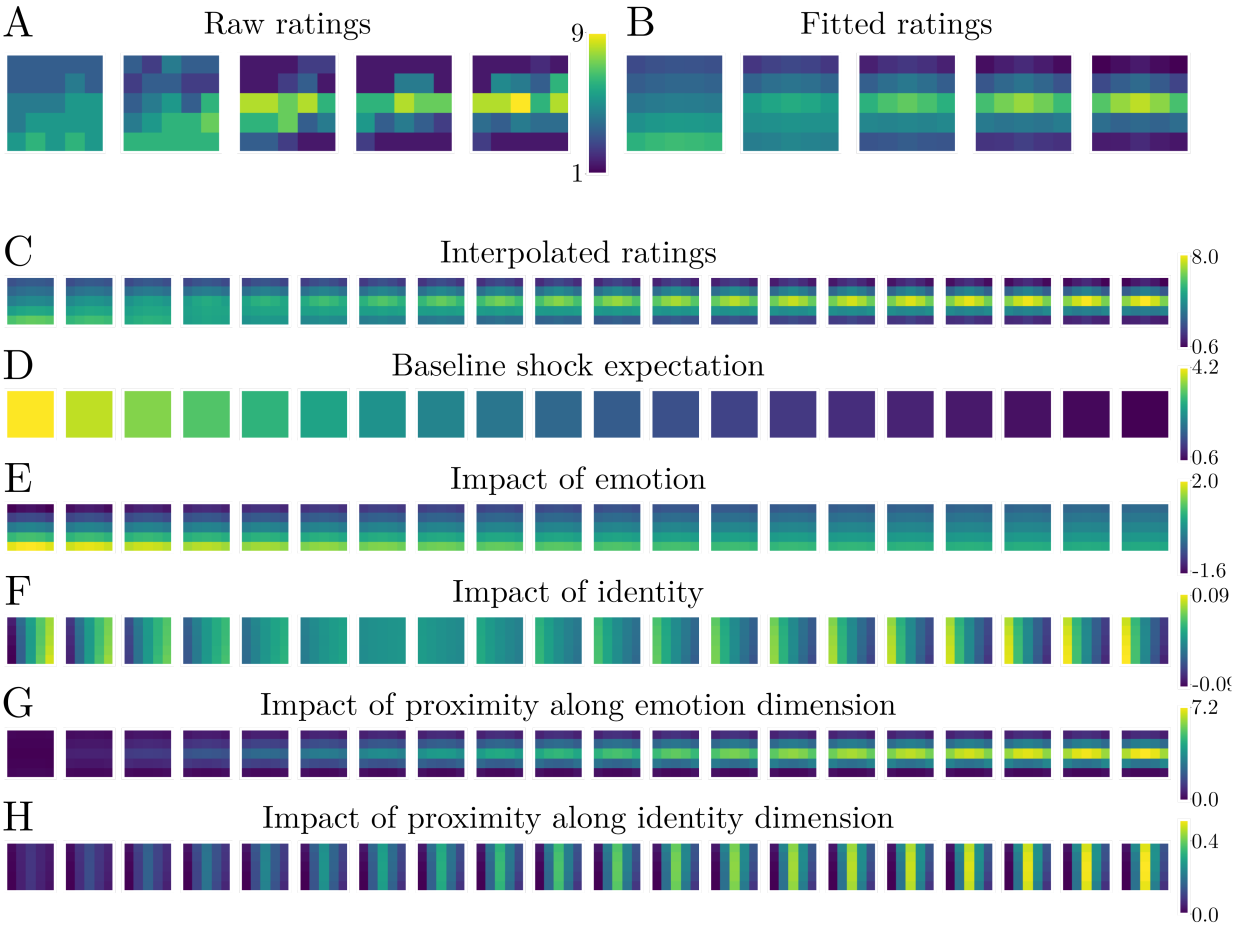


**Figure S2: Interpolation of shock expectations and features.** In order to include the modeled shock expectation and the constituent features as parametric modulators, we interpolated them from the 5 ratings to twenty microblocks. (**A**) Modeling the raw ratings at 5 time points yielded (**B**) modeled shock expectations. (**C**) Those were interpolated to 20 time points. The same procedure was applied to the constituent features, namely (**D**) the baseline shock expectation, (**E**) the impact of emotional expression, (**F**) the impact of identity, (**G**) the impact of proximity to the reinforced stimulus along the emotion dimension and (**H**) the impact of proximity to the reinforced stimulus along the identity dimension.

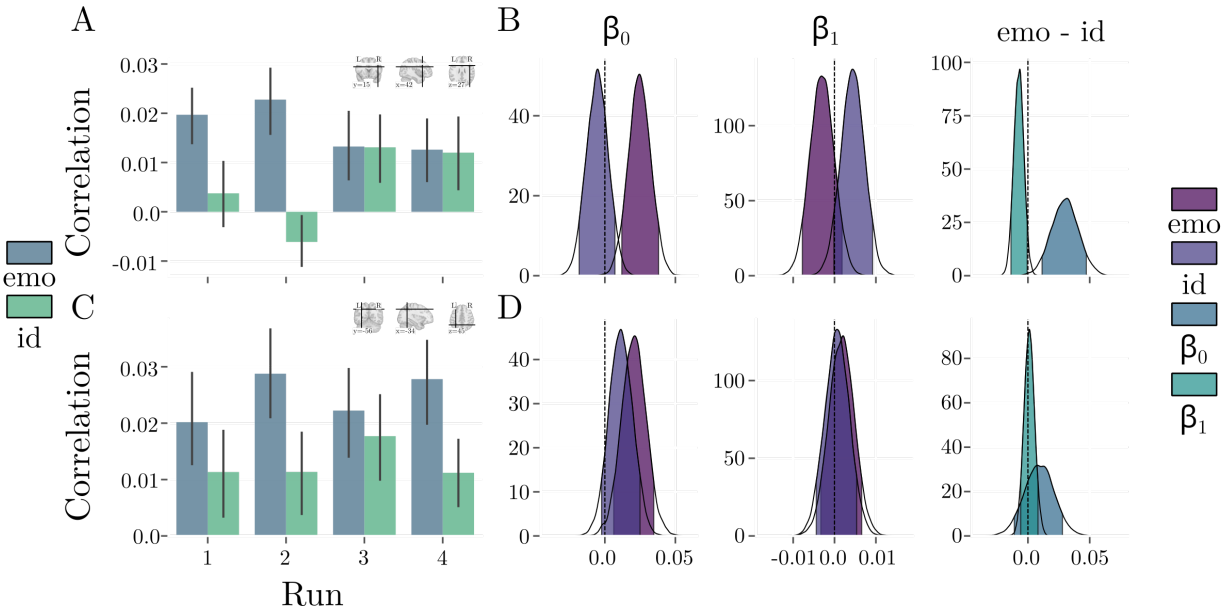


**Figure S3: Additional RSA results.** We repeated the Bayesian hierarchical linear regression with correlations from the right MFG (**A-B**) and the left IPS (**C-D**). (**A**) Correlations in the right MFG showed a similar pattern as in the left MFG. Representations initially only depended on the emotion dimension but on both dimensions in the second half of the experiment. This transition was less smooth than in the left MFG. (**B**) In line with that, posterior estimates favored the same direction of effects, but less decisively so. (**C**) In contrast, representations in the left IPS were more static and depended on both dimensions throughout the experiment, with more emphasis on the identity dimension. (**D**) This pattern was corroborated by posterior estimates, with positive intercepts but slopes that were close to zero.

| **Experiment** | **Condition** | **Model** | **ELPD** | **ΔELPD (SE)** |
|---|---|---|---|---|
| Experiment 1 | Angry | Fixed | -20472.03 | 0 (-) |
| | | Dynamic | -20562.04 | -90.01 (27.40) |
| | Happy | Fixed | -20164.23 | 0 (-) |
| | | Dynamic | -20240.10 | -75.86 (28.70) |
| Experiment 2 | Angry | Fixed | -19037.65 | 0 (-) |
| | | Dynamic | -19091.26 | -53.60 (28.27) |
| | Happy | Fixed | -18672.03 | 0 (-) |
| | | Dynamic | -18800.57 | -128.55 (28.53) |

**Table S1: Model comparison for perceptual spaces.** In the first two experiments, we compared two models for the perceptual spaces. One that assumes fixed perceptual spaces (Fixed) and one that assumes that perceptual spaces could be subject to change during the conditioning procedure (Dynamic). In both experiments and all conditions, the model assuming fixed perceptual spaces fit the data considerably better as indicated by the differences in expected log pointwise predictive density (ELPD) and the corresponding standard errors (SE).

| Experiment | Condition | Model | ELPD | ΔELPD (SE) |
|---|---|---|---|---|
| Experiment 1 | Angry | Partial DR | -11224.97 | 0 (-) |
| | | No DR | -11377.81 | -152.83 (25.82) |
| | | Full DR | -11451.33 | -226.36 (28.22) |
| | Happy | Partial DR | -10596.04 | 0 (-) |
| | | No DR | -10777.57 | -181.53 (33.30) |
| | | Full DR | -10827.97 | -231.93 (23.90) |
| Experiment 2 | Angry | Partial DR | -11562.07 | 0 (-) |
| | | No DR | -11737.08 | -175.02 (30.41) |
| | | Full DR | -11791.10 | -229.04 (22.94) |
| | Happy | Partial DR | -11717.04 | 0 (-) |
| | | No DR | -11915.10 | -198.06 (21.74) |
| | | Full DR | -11984.19 | -267.15 (36.62) |
| Experiment 3 | Angry | Partial DR | -11077.13 | 0 (-) |
| | | No DR | -11154.05 | -76.92 (26.53) |
| | | Full DR | -11409.12 | -332.00 (26.76) |
| | Happy | Partial DR | -12558.48 | 0 (-) |
| | | No DR | -12703.50 | -145.02 (17.47) |
| | | Full DR | -12802.83 | -244.36 (32.71) |

**Table S2: Model comparison for behavioral models.** We fit three different models to the behavioral data. These models differ in their assumption of dimensionality reduction. One model assumes a full reduction of the perceptual space to only the emotional dimension (Full DR), one model assumes a partial reduction of either dimension (Partial DR) and one model assumes no reduction at all (No DR). For a detailed description of the models, see "Methods". Model comparison indicated that the model assuming partial dimensionality reduction fit the data best in all experiments and conditions.
